# A phylogeny-based metric for estimating changes in transmissibility from recurrent mutations in SARS-CoV-2

**DOI:** 10.1101/2021.05.06.442903

**Authors:** Damien Richard, Liam P Shaw, Rob Lanfear, Russell Corbett-Detig, Angie Hinrichs, Jakob McBroome, Yatish Turakhia, Mislav Acman, Christopher J Owen, Cedric CS Tan, Lucy van Dorp, François Balloux

## Abstract

Severe acute respiratory syndrome coronavirus 2 (SARS-CoV-2) emerged in late 2019 and spread globally to cause the COVID-19 pandemic. Despite the constant accumulation of genetic variation in the SARS-CoV-2 population, there was little evidence for the emergence of significantly more transmissible lineages in the first half of 2020. Starting around November 2020, several more contagious and possibly more virulent ‘Variants of Concern’ (VoCs) were reported in various regions of the world. These VoCs share some mutations and deletions that haven arisen recurrently in distinct genetic backgrounds. Here, we build on our previous work modelling the association of mutations to SARS-CoV-2 transmissibility and characterise the contribution of individual recurrent mutations and deletions to estimated viral transmissibility. We then assess how patterns of estimated transmissibility in all SARS-CoV-2 clades have varied over the course of the COVID-19 pandemic by summing transmissibility estimates for all individual mutations carried by any sequenced genome analysed. Such an approach recovers the Delta variant (21A) as the most transmissible clade currently in circulation, followed by the Alpha variant (20I). By assessing transmissibility over the time of sampling, we observe a tendency for estimated transmissibility within clades to slightly decrease over time in most clades. Although subtle, this pattern is consistent with the expectation of a decay in transmissibility in mainly non-recombining lineages caused by the accumulation of weakly deleterious mutations. SARS-CoV-2 remains a highly transmissible pathogen, though such a trend could conceivably play a role in the turnover of different global viral clades observed over the pandemic so far.

**Caveats:**

- This work is not about the severity of disease. We do not analyse the severity of disease. We do not present any evidence that SARS-CoV-2 has decreased in severity.
- Lineage replacement dynamics are affected by many factors. The trend we recover for a decrease in inferred transmissibility of a clade over time is a small effect. We caution against over-interpretation. This result would not affect the management of the SARS-CoV-2 pandemic: for example, we make no claims about any impact on the efficacy of particular non-pharmaceutical interventions (NPIs).
- Our phylogeny-based method to infer changes in estimated transmissibility due to recurrent mutations and deletions makes a number of simplifying assumptions. These may not all be valid. The consistent trend for the slight decrease we report might be due to an as-yet-unidentified systematic bias.

## Introduction

Severe acute respiratory syndrome coronavirus 2 (SARS-CoV-2), the viral agent of the COVID-19 pandemic, has been acquiring genomic diversity since its emergence in humans in late 2019^1–3^. The unprecedented scale of generation and sharing of SARS-CoV-2 genomes has allowed tracking of the accumulation of mutations in close to real time; supporting epidemiological tracing of transmission, mutation surveillance and phylogenetic analyses^4–8^. Such analyses have highlighted a high rate of recurrent substitutions and deletions (homoplasies) in SARS-CoV-2 alignments^1,9–12^ and more recently allowed for the rapid flagging of ‘Variants of Interest’ (VoIs) and ‘Variants of Concern’ (VoCs) associated with higher transmissibility, immune evasion or higher virulence.

Mutations accumulate in SARS-CoV-2 at a rate of approximately two mutations per lineage per month^1,13^. Mutations arise following stochastic errors during replication or can be induced by host anti-viral editing proteins leading to characteristic mutational biases^9,14–17^. Mutations may also be exchanged between lineages through recombination events, combining genetic material from different viruses simultaneously infecting the same host into a new lineage^18^. While recombination is often presumed to be widespread in coronaviruses, at least within species, evidence in SARS-CoV-2 has remained scarce until recently, likely due to the limited power to detect genetic recombination provided by current levels of genetic diversity^19^ (https://observablehq.com/@spond/linkage-disequilibirum-in-sars-cov-2, accessed 11^th^ October 2021). Recent studies however provide compelling evidence that some recombinants are circulating at low incidence levels^20–24^. In addition to point mutations, the possible evolutionary importance of genomic deletions has come to the fore, in particular in the context of antigenicity^8,10,25,26^.

To date, all observed SARS-CoV-2 lineages have been highly genetically similar to one another (September 2021). Over the first ten months of the pandemic (December 2019 – September 2020) there was fairly limited evidence for marked variation in transmissibility or virulence between different SARS-CoV-2 lineages^9^. Patterns of emerging genomic diversity were well explained by neutral evolutionary processes, with the exception of the D614G mutation which has been associated with higher transmissibility^27,28^. However, towards the end of 2020, several lineages of SARS-CoV-2 attracted attention following their rapid increase in frequency in regions where they were first observed and the constellations of mutations they harbour^29^. Those include Alpha (also known as 501Y.v1 or PANGO lineage B.1.1.7 or NextStrain clade 20I/501Y.v1) first detected in the UK^30^, Beta (a.k.a. 501Y.v2 or PANGO lineage B.1.351; Nextstrain clade 20H/501Y.v2) first detected in South Africa^31^, Gamma (a.k.a. 501Y.v3 or PANGO lineage P.1; Nextstrain clade 20J/501Y.v3) first detected in cases linked to Brazil^32^ and Delta (a.k.a. Nextstrain clade 21A including the PANGO lineage B.1.617.2 and its daughter AY lineages) first detected in India in late 2020^33^.

Notable features of the VoCs are that they carry multiple mutations within the spike protein, a region crucial for human receptor binding and a dominant target for neutralizing antibodies during infection^34,35^. These include the 501Y mutation observed in Alpha, Beta and Gamma, 484K which independently occurred in Beta and Gamma and in some cases Alpha; 452R (observed in Delta), the spike codon 417 (417N in Beta and 417T in Gamma) and the spike codon 681 (681H in Alpha and 681R in Delta). Such point mutations may be coupled with deletions in the spike N-terminal domain (NTD) which have been observed repeatedly, including during chronic infections^8,36–38^, with H69 V70 (Δ69-70) and Y144 (Δ144) found together in Alpha, Δ243-244 found in Beta and Δ156-157 in Delta. All three VoCs carrying 501Y also have the same three amino acid deletion in NSP6 (Δ106-108). Interestingly, while many of the mutations present in VoCs were observed far earlier in the course of the pandemic^7^, they seemingly conferred no easily detectable adaptive advantage until more recently, pointing to a possible shift in the SARS-CoV-2 landscape of selective pressures and/or epistasis^9,12^.

Each of the VoCs have measurable phenotypic effects compared to the Wuhan-Hu-1 reference genome (the original ‘wild-type’ representative), including enhanced receptor binding in each case^39–41^, increased transmissibility^32,33,42,43^ and some ability to evade past immunity from natural infection and/or vaccination for Beta, Gamma and Delta^33,44–49^. Real-world effects on mortality require carefully controlling for multiple factors, but infection with Alpha has been associated with higher hospitalisation rates by several studies, even if mortality in hospitalised patients seems unaffected^50–52^. Higher virulence of Gamma^53^ has also been suggested^53^. A further increase in virulence of the Delta variant over Alpha has been reported^54,55^, which may stem from a pleiotropic effect of their increased infectivity.

All VoCs exhibit an excess of non-synonymous mutations (i.e., dN/dS>1) consistent with adaptive evolution^30–32,56^. It has been suggested, based on both the large number and nature of its mutations, that Alpha spread into community transmission following a period of rapid evolution in a chronically infected patient^12,30^. An alternative hypothesis is that the combination of mutations observed in Alpha may have been generated through recombination between circulating SARS-CoV-2 lineages, although phylogenetic tests of recombination have not found evidence of a recombinant origin of the alpha strain^20–24^. Emergence through the rapid accumulation of mutations during a chronic infection seems an unlikely scenario for the origin of the other three VoCs, insofar basal strains carrying only a subset of the constellation of VOC-defining mutations have been characterised. Irrespective of how the different VoCs emerged, the occurrence of a number of common recurrent mutations and deletions suggests some action of convergent evolution, likely driven by the phenotypic advantage of increased human ACE2 receptor binding affinity and/or some ability to bypass prior immunity^12^.

We previously developed a phylogenetic index to identify all recurrent mutations in global SARS-CoV-2 phylogenies and tested their association to variation in estimated transmissibility^9^. When applied to genome assemblies shared over the first ∼seven months of the pandemic (up to 30/7/2020), our method did not identify any recurrent mutation that had a statistically significant association with increased estimated transmissibility on its own. The multiple emergences of more transmissible VoCs in late 2020, which also involve recurrent deletions, motivated us to extend our phylogenetic scoring framework. Here, we modify our phylogeny-based method to include notable recurrent deletions and analyse a far larger dataset of over half a million globally distributed SARS-CoV-2 genome assemblies released via the GISAID Audacity platform^5,6^. We characterise the distribution and number of emergences of all mutations and deletions over the phylogeny (as of 29/08/2021), before filtering for a well-supported set of recurrent mutations to test for their association with estimated transmissibility. We express the relative contribution to estimated transmissibility provided by carriage of any individual recurrent mutation or selected deletion as Coefficients of Exponential Growth Alteration (CEGAs). Finally, we show that a simple genetic model, combining the transmissibility effect of each individual CEGA into a multilocus per-isolate score, can largely recover expected relationships in relative estimated transmissibility of major SARS-CoV-2 clades from previous work. This suggests that we can use this phylogeny-based metric to evaluate relative changes in the estimated transmissibility of viral clades over time.

While our approach uncovers a marked increase in estimated transmissibility of SARS-CoV-2 clades in circulation following the global spread of the Alpha VoC, and a second one following the spread of the Delta VoC, we also find evidence that the estimated transmissibility within major clades tends to subtly decrease over time since their first detection. Such an observation could be explained by the accumulation of weakly deleterious mutations in lineages undergoing limited genetic recombination, which is a well-known evolutionary phenomenon^57,58^.

## Results

### SARS-CoV-2 global genetic diversity

We consider the global genetic diversity of SARS-CoV-2 assemblies generated from samples collected between the 26^th^ of December 2019 to the 20^th^ of August 2021 (Supplementary Figure S1) released via the GISAID Audacity platform^5,6^. These encompass 3,650 locations spanning six continental regions and comprise 126,356, 2,150, 13,986 and 255,774 from each of the Alpha, Beta, Gamma and Delta VoCs, respectively. In contrast to the diffuse geographic structure observed early in the pandemic, when different SARS-CoV-2 lineages were essentially randomly distributed globally due to multiple introduction events in various countries^1,59^, stronger patterns of geographic structure have now emerged (Supplementary Figure S1a). However, SARS-CoV-2 genetic diversity remains low at this stage and the majority of currently circulating lineages are represented by intermediate basal isolates starting from the root. Notable exceptions are a few dominant lineages located on longer branches away from the root, possibly suggestive of a burst of adaptive mutations^30^. The large number of assemblies included in this dataset means that (i) all of the 29,903 nucleotide positions of the Wuhan-Hu-1 reference genome carry single nucleotide variants (SNVs) in at least one sample, (ii) a large fraction (>99%) of SNVs independently emerged multiple times, and (iii) multiple alternate alleles are often present at a single nucleotide position, which poses a challenge for present-day bioinformatics approaches.

### Recurrent substitutions and deletions in SARS-CoV-2

Across 491,449 SARS-CoV-2 genomes we detect 1,467 SNVs at a frequency of at least 0.1% (Figure 1, Figure S1). Most (>99%) of these can be considered homoplastic, appearing to have emerged independently more than once given the global SARS-CoV-2 phylogeny. Phylogenetic uncertainty is expected to lead to an overestimation of the number of homoplasies. This can occur because most SARS-CoV-2 genomes contain relatively little phylogenetic information, meaning that clades of closely related genomes can be interspersed due to phylogenetic noise induced by errors in genome sequencing or assembly. This interspersion is known to elevate estimates of homoplasy^60^, so herein we focus on mutations that have arisen a very large number of times in the phylogeny in genetically distinct SARS-CoV-2 clades. Consistent with previous observations^9,16^, a large fraction of the 1,467 homoplasies we detect (48%) derive from C→T changes likely caused by host anti-viral RNA editing machinery^14,15,17^ (Supplementary Table S2, Supplementary Figure S2). We note that the minimum numbers of emergences estimated for synonymous and non-synonymous recurrent mutations were significantly different largely driven by observations over ORF1ab, ORF7a and the nucleocapsid (Supplementary Figure S3).

**Figure 1.**
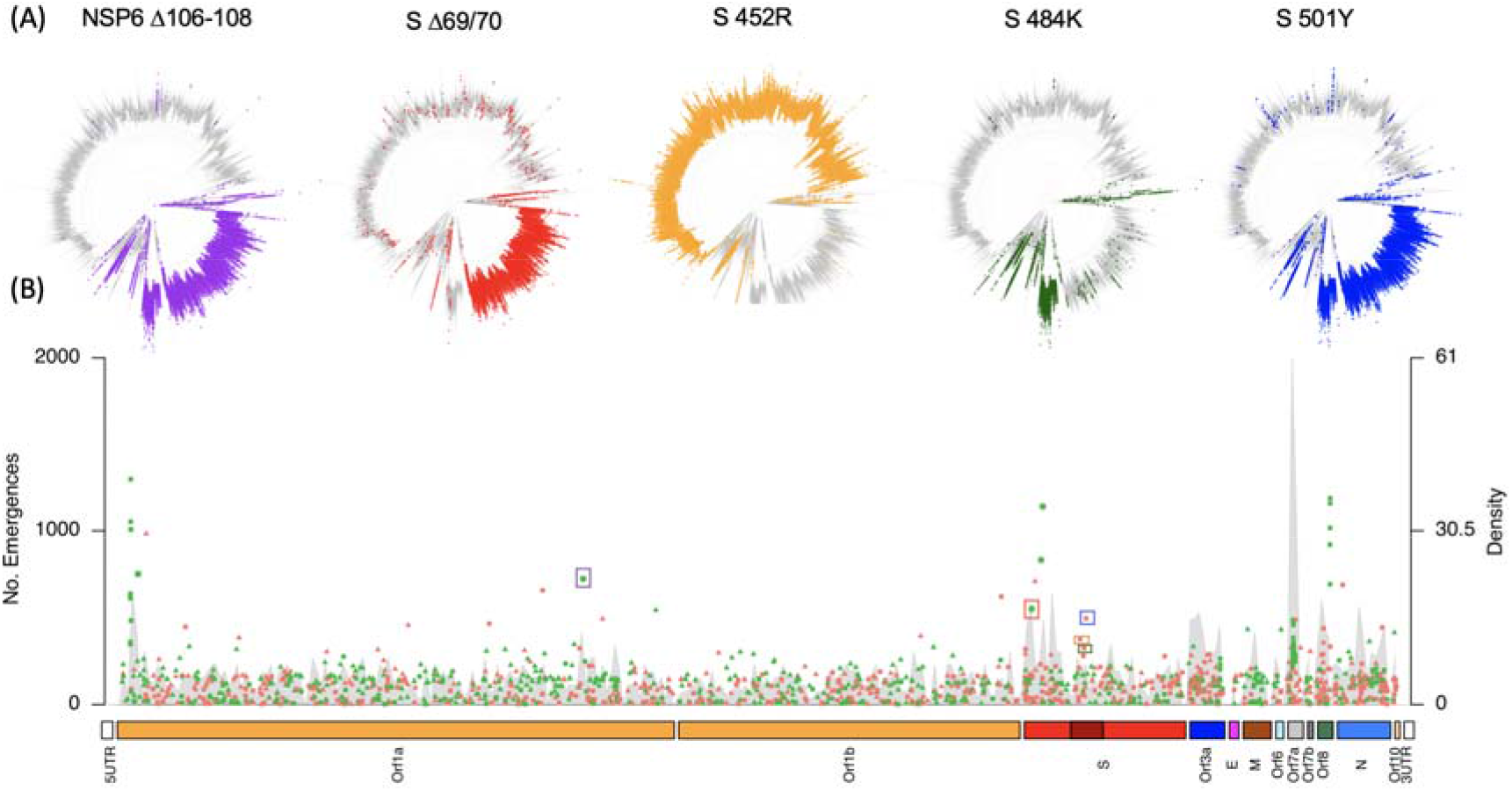
(A) Placement of unfiltered homoplasies associated to some VoCs in the global phylogeny; NSP6 Δ106-108 (nucleotide positions 11288-11296; 724 emergences since 10/03/2020), Spike (S) Δ69/70 (nucleotide positions 21765-21770; 550 emergences since 14/04/2020), Spike 452R (nucleotide position 22,917; 373 emergences since 21/04/2020), Spike 484K (nucleotide position 23,012; 321 emergences since 10/03/2020) and Spike 501Y (nucleotide position 23,063; 494 emergences since 14/04/2020). (B) Bottom panel provides the number of emergences of all recurrent mutations detected along the SARS-CoV-2 genome, with sites depicted in the phylogenies highlighted by coloured boxes, together with the density of homoplasies over an 80-nucleotide sliding window (y-axis at right). Synonymous changes are shown in green, non-synonymous changes are shown in orange. Those sites which correspond to C→T or T→C changes are shown with a triangle. Studied deletions are shown with a ‘*’. All other mutations are shown with a square.

Genome-wide we identify several mutations that have emerged a large number of times, in particular in the spike protein and ORF8 (Figure 1, Supplementary Figure S4). Mutations commonly associated with VoCs have appeared in multiple genetic backgrounds, already well before the first reports of VoCs towards the end of 2020 (Figure 1). For instance, spike deletion Δ69/70, present in Alpha and several other PANGO lineages, was first detected in our dataset on 14/04/2020. Since then we estimate it has emerged independently a minimum of 550 times in the phylogeny, including in combination with other mutations in the spike protein such as N439K and Y453F^10,61^. We estimate a minimum of 321 emergences of E484K (earliest observation 10/03/2020), 373 emergences of L452R (earliest observation 21/04/2020) and 494 emergences of N501Y (earliest observation 14/04/2020). While phylogenetic misplacement will affect these raw estimates (see Methods and above), they nonetheless highlight the long duration of circulation of major mutations of interest in VoCs and their propensity to independently arise on the background of many different SARS-CoV-2 lineages.

### Effect of mutations and deletions on SARS-CoV-2 estimated transmissibility

By focusing our observations on mutations and deletions which have evolved independently multiple times, often in different locations and epidemiological settings, we largely exclude consideration of those sites which may have risen to high frequency solely due to founder effects^1,9,62^. To test across observations for association between individual recurrent mutations or deletions and variation in SARS-CoV-2 estimated transmissibility, we apply a phylogenetic scoring approach. The model we develop assumes that a transmission advantage conferred by a mutation/deletion translates into an excess of descendants in the phylogeny, given random sequencing of isolates. We can quantify this effect by considering the relative number of descendants that the lineage carrying the mutation/deletion gives rise to relative to its sister lineage in the phylogeny which lacks the mutation/deletion^9^ (Supplementary Figure S5). To do so, we study pairs of SARS-CoV-2 lineages in the phylogeny which descend from nodes corresponding to the hypothetical ancestor which acquired a particular mutation/deletion. We assess these ratios of descendants across independent emergences of the mutation/deletion in the phylogeny (homoplastic replicate). To express these ratios in terms of selection differentials (*s*), we then normalise by the estimated number of viral generations over which the lineages have been observed in circulation (see Methods). We refer to this normalised index as Coefficients of Exponential Growth Alteration (CEGA). CEGA provide estimates of the change in transmissibility due to a focal mutation. Hence an average CEGA of zero suggests that a mutation has no effect on transmissibility. A CEGA greater than zero suggests that the focal mutation increases the estimated transmissibility, and vice-versa for a CEGA score below zero.

As noted, recurrent mutations are to be expected within SARS-CoV-2 phylogenies given the well documented role of immune-mediated hyper-mutation in introducing genomic diversity ^9,14–17^ and may also be observed due to a degree of phylogenetic misplacement^63^. We therefore implement a series of filtering steps so that we only assess, using the CEGA metric, phylogenetically well-supported recurrent mutations and deletions. These steps include: (i) only taking forward for analysis mutations for which we detect at least five independent emergences (replicates) with (ii) sufficient number of descending offspring displaying each allele (iii) excluding erroneous positions associated to sequencing artefacts and (iv) restricting our analysis to nodes defining sister lineages with high allele frequencies (see Methods). In this way we exclude from consideration low frequency and singleton emergences which are more likely to have arisen due to sequencing errors or phylogenetic misplacement^1,11,16^.

Of the 1,467 identified homoplasies, 719 (50%) passed all selection criteria for downstream analysis. These 719 sites comprise 713 substitutions and six deletions (Supplementary Table S3). On average each of the 719 recurrent mutations was represented by 10.4 ± 7.6 (mean ± SD) independent emergences on the phylogeny that passed our filters, though with considerable variance. The number of well supported emergences by nucleotide position is provided in Supplementary Table S3. Of these 719 sites, 155 displayed a positive CEGA score (i.e., associated to on average higher estimated transmissibility). Values ranged from −0.34 to 0.32 across sites with a slightly negative mean (−0.044) and median (−0.048). Consistently, we observe a significant deviation from a normal distribution (Shapiro-Wilk *p*-value of <1×10^−9^ in each case) to a right-tailed distribution (mean skewness 0.49) suggesting that the majority of recurrent mutations/deletions in SARS-CoV-2 have a slightly deleterious effect on estimated transmissibility. We obtain highly similar distributions for the mean and median CEGA score per site (Figure 2, Supplementary Figure S6-S8) suggesting averaging CEGA scores of individual mutations/deletions can robustly capture their associations to estimated transmissibility in multiple genetic backgrounds (Figure 2, Figure 3A).

**Figure 2:**
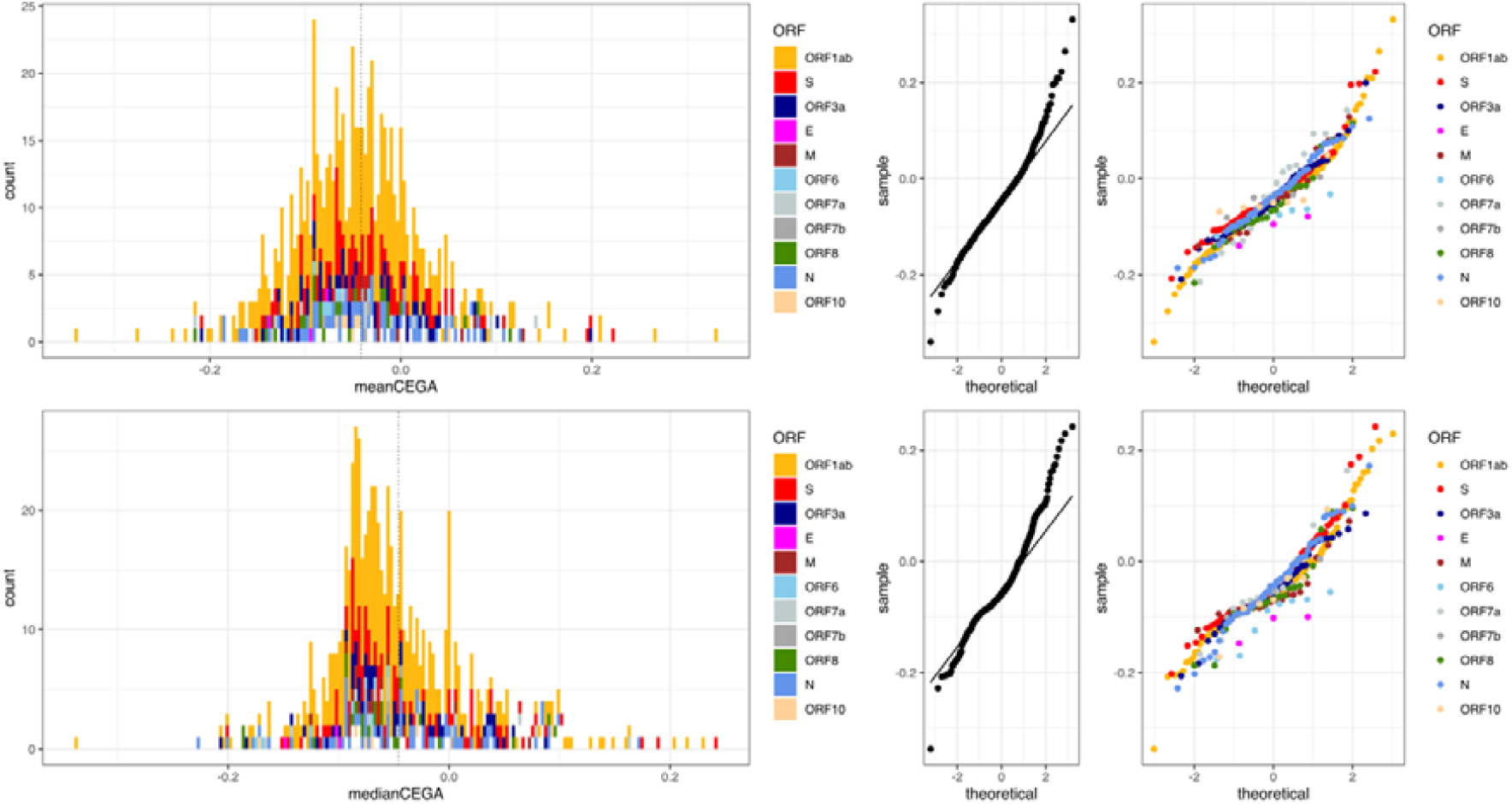
Distribution of mean (top) and median (bottom) CEGA score for 719 homoplastic mutations which passed our scoring filters. In both cases the mean values of both estimates fell below 0 (−0.044 for mean CEGA; −0.048 for median CEGA). Right hand panels provide the qqplots for included sites genome-wide and coloured by genomic feature.

**Figure 3:**
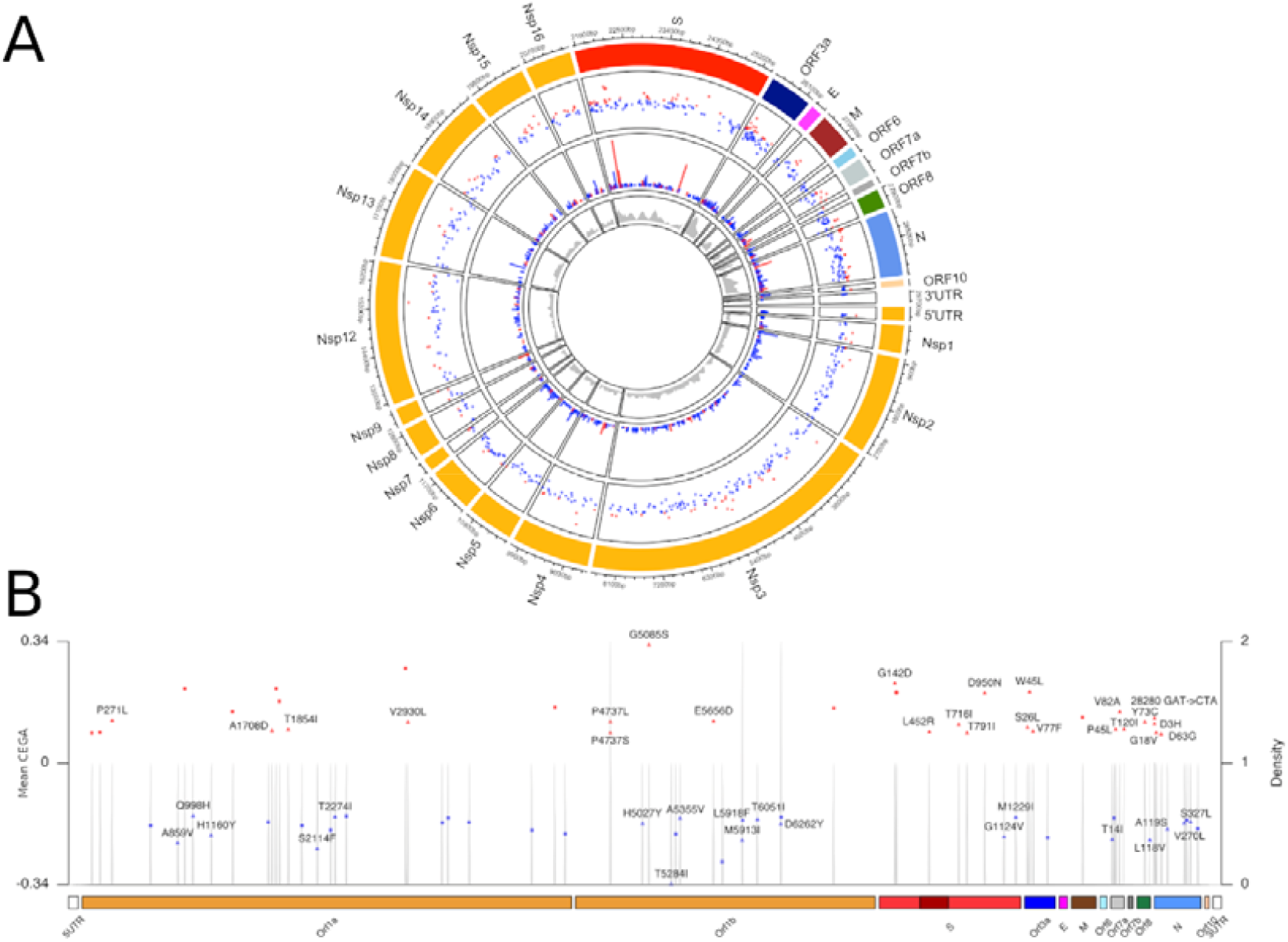
(A) Circular representation of the genomic structure of SARS-CoV-2 with associated mean CEGA scores. From outer to inner circles: Gene names; mean CEGA score (red: positive; blue: negative) with those estimated at deletions denoted *; Number of independent emergences used for CEGA computation; density of sites tested using a window of 20 nucleotides. (B) Sites along the genome with mean CEGA scores plotted for those falling in the upper and lower 5% of estimates. Non-synonymous sites are highlighted with a triangle with associated amino acid change, synonymous sites are depicted with a square with associated nucleotide change, deletions are shown with a *. Grey sharing provides the density of high/low scoring sites over a 20-nucleotide sliding window. As in (A) positive scores are depicted red with negative scores depicted blue. Figure produced using the R package circlize 0.4.12 ^70^ and karyoploteR 1.16.0 ^71^.

CEGA scores associated with C→T or G→T transitions tended to fall below zero on average (p<2.2e-13), suggesting an excess of such sites leads to a slightly deleterious effect on estimated transmissibility (Supplementary Figure S8). However, we otherwise recorded no strong trends associating any particular type of mutation to higher or lower CEGA scores (Supplementary Figures S7-S8). Considering those sites leading to the most extreme CEGA scores - 35 mutations within the upper 5% of mean CEGA scores - nine correspond to synonymous changes, 24 to non-synonymous changes and two to deletions (Δ119/120 in ORF8 and Δ156/157 in the spike, both found in the Delta VoC). Conversely, of the 35 mutations associated with the lowest 5% of CEGA estimates, 16 corresponded to synonymous changes and 19 to non-synonymous changes, 11 of which are C→T transitions (Figure 3B). One of the most negative scores (CEGA=−0.21) observed is a non-synonymous change in the SARS-CoV-2 spike protein A1124V, highlighting the potential for changes in the spike protein to contribute to both higher and lower estimated transmissibility. Consistently, there was no overall tendency for recurrent mutations in the spike protein to give rise to significantly higher or lower CEGA scores than those obtained in other structural genes (Supplementary Figure S9). Of our curated set of six deletions, three: spike Δ144; spike Δ156/157 and ORF8 Δ119/120 were associated with positive CEGA values, and NSP1 Δ141- 143; NSP6 Δ106-108 to a negative value (Supplementary Table S3). Interestingly, the spike Δ69/70 deletion found in the Alpha variant was associated to an overall negative score, but a positive value (0.094) when restricting the analysis to the five homoplasies observed within Alpha, suggesting that its effect on transmissibility may have some dependence on the genetic background on which it is found.

Interestingly, the E484K mutation present in the spike protein of Beta and Gamma has an associated mean CEGA score of −0.065 suggesting it slightly reduces estimated transmissibility on average. Since, E484K is known to reduce convalescent serum neutralisation^26,64^ and implicated in reinfections^65^, this may point to an evolutionary trade-off between transmissibility and the propensity to (re-)infect immunized hosts. Among the VoCs, Delta stands out as it carries eight of the 35 highest 5% CEGA scores (Figure 3B), including seven non-synonymous mutations (spike: 3; N protein: 1; ORF3a: 1; ORF7a: 2). We also estimate three high scoring mutations on each one of the three nucleotides encoding for nucleocapsid D3 amino-acid (28280:GAT->CTA)^66^. This triple mutation, leading to D3L, is predominantly found in Alpha, and has been implicated in greater subgenomic RNA expression in Alpha^67^ as a possible contributor to the higher transmissibility of this VoC. Also, P681R, close to the furin cleavage site^68^ and one of the signature mutations of Delta claimed to be responsible for its transmissibility increase over Alpha (which carries P681H)^69^, showed a positive CEGA score (CEGA=0.054).

Four out of 15 tested recurrent mutations located in the Receptor Binding Domain (RBD) of the spike protein (nucleotide positions 22,559-23,143^72^) displayed positive CEGA values, in increasing order of value: S494P (CEGA=0.013), T478K (CEGA=0.027; present in Delta), S477N (CEGA=0.046, associated with 20A.EU2) and L452R (CEGA=0.087; present in Delta). While we did not recover any significant correlation between the estimates for seven of these RBD recurrent mutations and ACE2 receptor binding affinity measured following a deep mutational scan analysis^39^, we do detect a significant correlation between mean CEGA values and RBD expression (adj R^2^=0.6, *p*<0.001) (Supplementary Figure S10).

### Evaluation of the transmissibility of SARS-CoV-2 clades through time

We next sought to ask whether it is possible to combine individual CEGA scores to estimate the relative transmissibility of a SARS-CoV-2 isolate from its genome. There are many ways to potentially combine CEGA scores into a single genome-wide estimate. The best way to do so likely depends on the underlying (and unknown) genetic architecture of transmissibility. Regardless, it is clear that if a method of combining CEGA scores were successful, it should at least allow one to recover the relative estimated transmissibility of any genome in the phylogeny, because this is precisely the data that was used in the estimation of individual CEGA scores. Such a combined score, or ‘genome-wide transmissibility coefficient’, offers a quantitative estimate of changes in estimated transmissibility, under the assumption of no epistatic interactions between any of the mutations and deletions considered.

To estimate a multi-locus CEGA score we considered all recurrent mutations and deletions which passed our filters, then combined them in a multiplicative fashion into a ‘poly-CEGA’ score for each genome (see Methods). Initial inspection of the poly-CEGA scores over a representative sample of the dataset highlights that the transmissibility of SARS-CoV-2 has altered in a stepwise manner over the COVID-19 pandemic so far (Figure 4), with two marked increases in the poly-CEGA scores following the global spread of the Alpha and Delta VoCs.

**Figure 4.**
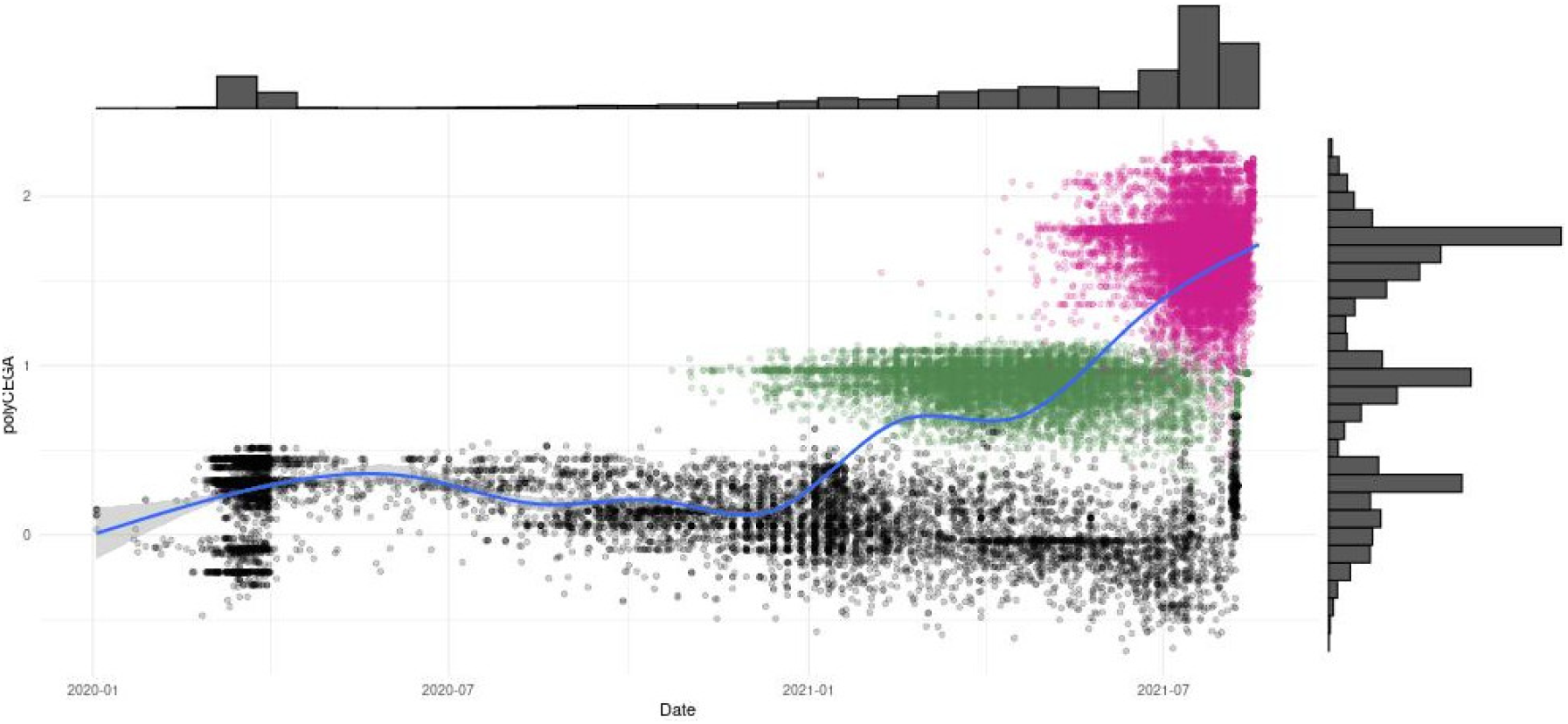
Poly-CEGA values (y-axis) of all available assemblies from December 2019 to April 2020 and a random subset from April 2020 onwards (x-axis). Individual genomes (points) coloured in green label the Alpha VoC, those coloured in pink label the Delta VoC. All other considered genomes are coloured in black.

To assess whether the poly-CEGA framework recovers relative differences in estimated transmissibility between SARS-CoV-2 clades, we computed poly-CEGA estimates for genomes assigned to different clades defined by NextStrain^4^, encompassing all current VoCs (https://clades.nextstrain.org/, accessed 10 September 2021). Calculating poly-CEGA scores over all major clades, we found the majority (10 out of 13 testable clades) have an average poly-CEGA score above 0. Surprisingly, Beta and Gamma showed negative polyCEGA values. By far the highest estimates were for Delta (mean poly-CEGA=1.85) and Alpha (mean poly-CEGA=0.93), which exhibited poly-CEGA scores significantly higher than all other considered SARS-CoV-2 clades (Wilcoxon *p*<1×10^−6^) (Figure 4, Supplementary Figures S11-S24). Of the mutations contributing the most to the high polyCEGA values recovered for both Alpha and Delta, we find the two mutations ORF1ab 3037 and NSP12 14408 which are associated to the D614G haplotype. The high polyCEGA score for Alpha also stems from the nucleocapsid triple mutation D3L and ORF8 Y73C, and that of Delta from the spike mutations G142D, L452R, P681R and D950N, but also spike deletion Δ156-157 and ORF7a V82A. We didn’t find any consistent pattern of gene contribution to polyCEGA across clades. Indeed, spike mutations are not necessarily the main drivers of high polyCEGA and are even sometimes driving low scores (Supplementary Figure S25).

These results suggest that the poly-CEGA scores can uncover relative variation in transmissibility among major lineages, as recovered using alternative methods such as methods that estimate relative transmissibility in different epidemiological settings^73^. We note, however, that our method requires observations (emergences) of mutations present in a clade of interest to fully capture relative patterns of transmissibility. Indeed, when we computed the CEGA scores using two restricted datasets comprising no assemblies assigned to Alpha or Delta respectively, we did not observe higher polyCEGA transmissibility scores for Alpha and Delta relative to other clades (Supplementary Figures S26 and S27). The latter result suggests that the association of some mutations with transmissibility is strongly lineage-specific, which would be in line with the moderate amount of convergent evolution observed between SARS-CoV-2 lineages until now.

Having explored patterns in estimated transmissibility over the global dataset (Figure 4) we next asked whether there are notable trends in poly-CEGA scores within major phylogenetic clades (Figure 5). Within defined SARS-CoV-2 clades, application of a simple linear model (poly-CEGA estimates against sampling time) reveals that in all cases aside from 19A, 20H and Delta (for which polyCEGA do not decrease over time), we recover a tendency for the average poly-CEGA score to subtly decrease with time (Supplementary Table S4). The relative size of this effect compared to the ‘initial’ increased transmissibility of a lineage is small but statistically significant (Figure 5C). Considering the ratio of positive to negative individual CEGA scores, in 11 out of 13 testable clades we identify a tendency towards accumulation of mutation which are associated to reduced estimated transmissibility, leading to a slight decrease in overall scores over time (Supplementary Figures S11-S24). Such an observation is consistent with the accumulation of weakly deleterious mutations over time.

**Figure 5.**
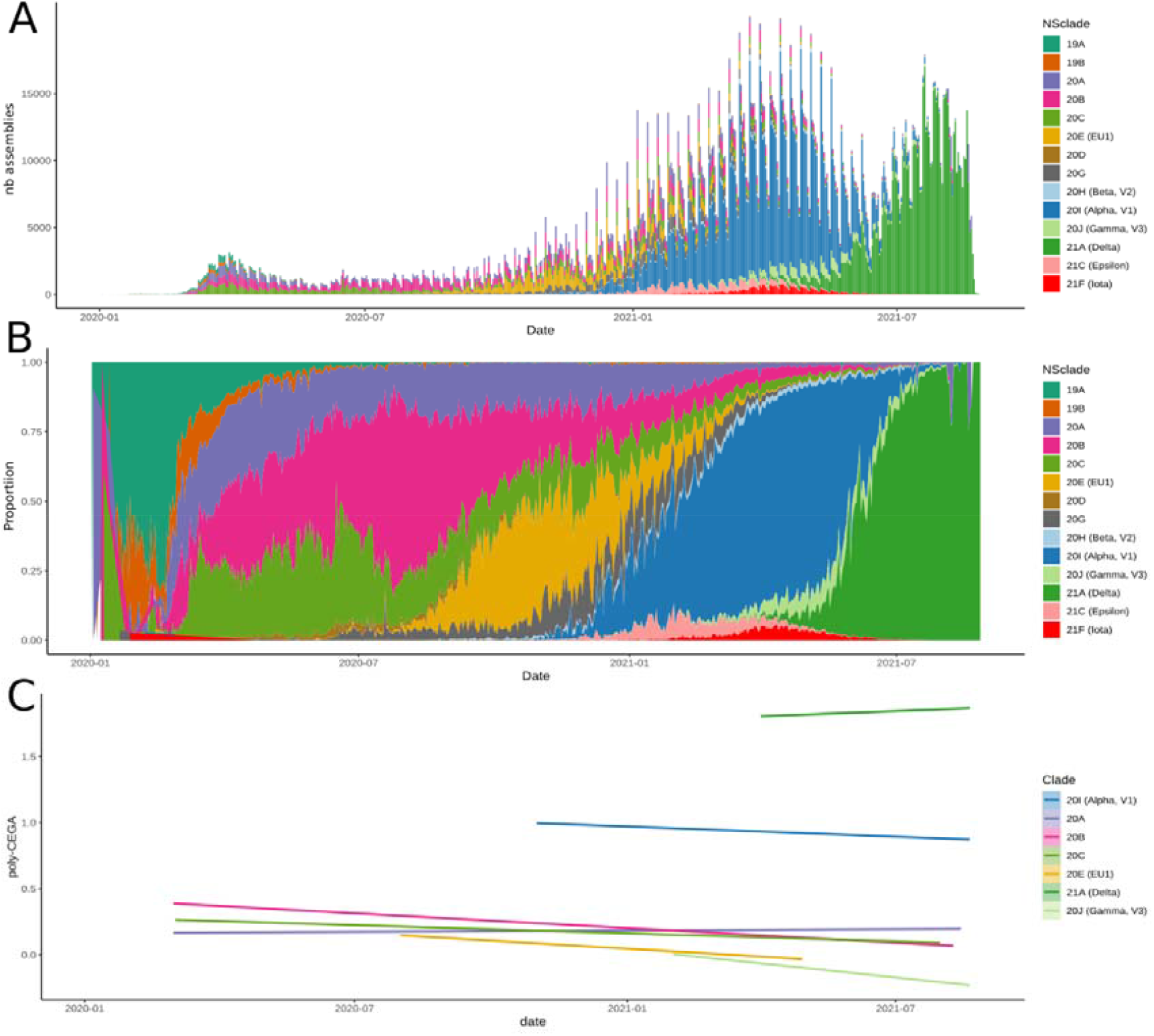
SARS-CoV-2 clades, defined by NextStrain, for genome assemblies available over the course of the pandemic dating from late December 2019 through to August 2021. (A) Histogram provides the daily number of sequenced SARS-CoV-2 samples from the GISAID database coloured by NextStrain clade, as given by the legend at right. (B) Daily frequency of each NextStrain clade estimated as the proportion of uploads to the GISAID genome database. (C) Linear regressions providing the temporal evolution of the poly-CEGA score – y axis (multiplicative assessment of the transmissibility of a genome from the mutations and deletions it carries) obtained for all genomes of each of the eight NextStrain clade for which more than 10,000 genome assemblies have been shared. The underlying values, per genome, are visualised in Supplementary Figures S11-S24 with values obtained from linear regression provided in Table S4. All three panels use the same colour code corresponding to the NextStrain clade assignment.

## Discussion

During the early stages of the COVID-19 pandemic, the evolution of SARS-CoV-2 followed a largely neutral dynamic. Apart from the rapid emergence and diffusion of the D614G haplotype^28^, little evidence for the emergence of viral lineages differing in their transmissibility or virulence was found^9,12^. This period of relative evolutionary stasis came to an end towards late 2020 with the detection of SARS-CoV-2 Variants of Concern (VoCs). The four VoCs (Alpha-Delta), each of which emerged independently in a different region, share some mutations and deletions. Such recurrently emerging changes are strong candidates for convergent evolution towards altered transmissibility.

In this work, we have estimated the number of emergences of more than 1,400 recurrent mutations and deletions detected in around half a million SARS-CoV-2 genomes sampled between December 2019 through to August 2021. Focusing on a well-characterised subset of these recurrent mutations and deletions, we assessed their association to the estimated transmissibility of SARS-CoV-2. We applied a phylogeny-based scoring metric which measures the relative number of progeny descending from nodes harbouring a specific, newly acquired change. The Coefficients of Exponential Growth Alteration (CEGA) scoring index provides estimates of the associations between all high frequency recurrent mutations and estimated transmissibility. It captures key components of SARS-CoV-2 evolution, including the dominance of C→T changes and the weakly deleterious overall effect of many mutations, in part driven by synonymous C→T and G→T changes.

While the effect of mutation carriage on estimated transmissibility appears modest for the majority of our tested sites, we identified a subset of recurrent changes giving rise to highly positive scores. While some of these have known phenotypic effects, this was not ubiquitously the case, for example many high scoring mutations fall outside of the SARS-CoV-2 spike protein. This suggests our approach holds value to identify sites for further functional characterisations and to support rapid assessment of combinations of mutations as they may appear in different backgrounds. Indeed, we found that when estimating the transmissibility of any SARS-CoV-2 genome based on its full genomic makeup, poly-CEGA scores performed well in recovering key aspects of the SARS-CoV-2 phylogeny, such as the global increase in estimated transmissibility following the emergence and the successive spread of the Alpha and later Delta VoCs (Figure 4).

Interestingly, when the transmissibility effect of individual mutations and deletions (CEGAs) is re-estimated without inclusion of any representatives of Alpha or Delta in the dataset, the resulting transmissibility scores of the genomes from those clades (polyCEGAs) does not recover higher transmissibility. This effect partly stems from our method: many of the mutations associated to high CEGA scores present predominately in Alpha and/or Delta do not pass our specified filters for computing robust individual CEGA scores in the downsampled datasets (Alpha: 26% and Delta: 53% of the total dataset), hence their contributions to the polyCEGA cannot be captured once strains from those lineages have been omitted from the analysis. Though, we find evidence for some mutations being associated to increased transmissibility in the genetic backgrounds where they are commonly found (epistasis). For example, the Δ69-70 deletion in the spike protein, which is found throughout the SARS-CoV-2 population was found to be associated to increased transmissibility in the Alpha variant, but not in other observed lineages. Lineage-specific effects of particular mutations on transmissibility is also compatible with the moderate overlap of mutations and deletions found at high-frequency in the Alpha and Delta variants, the only two variants which undeniably display higher transmissibility than previous and co-circulating clades so far.

Our work points to diverse genetic contributors to viral transmissibility. At this stage, we can only speculate on the possible functional mechanisms underlying enhanced transmission. One hypothesis relates to the relevance of mutations in the receptor binding domain that modify the biochemical phenotype of the virus^39^. While our results did not show evidence of a clear relation between the binding affinity effect of mutations and their estimated transmissibility, we did recover a significant relationship between the expression level of the receptor binding domain of a mutation and its CEGA score, although the relation between those phenotypes and viral fitness is complex^39^. An alternative, non-mutually exclusive mechanism for greater infective potential is the ability to escape neutralising antibodies primed by prior natural infection or vaccination, a situation which likely holds true for spike mutations at S477N, E484K and K417N/K417T^46,74–77^. An increase in transmissibility conveyed by the ability to bypass host immunisation will depend on the rate of vaccination and prior infection of the host population. Thus, contrary to mutations providing an intrinsic transmission advantage, those allowing the virus to bypass host immunisation may only be selected in host populations with significant immunity. Such variable selective pressure may contribute to the intriguing patterns we observe. For example, while the E484K and the K417N mutations have emerged many times, our results do not indicate that they contributed to an on average increase in estimated transmissibility (CEGA^E484K^ =-0.065; CEGA^K417N^ =-0.019).

Our results also highlight the importance of looking beyond the RBD of the spike protein for putative adaptive and recurrent changes, similar to related studies^73^. Indeed, we observe the greatest density of recurrent mutations within ORF7a, ORF3a, and the NTD of the spike protein (Figure 1); the latter being a highly diverse genomic region across Sarbecoviruses^78^ and a hotspot of antigenic evolution in human endemic coronaviruses^79,80^. A further intriguing outcome of extensive genomic surveillance efforts of SARS-CoV-2 is the marked patterns of lineage dynamics. Many clades are now essentially extinct, with others having been through fluctuations or rising/falling rapidly in frequency (Figure 5A-B). Our poly-CEGA estimates across SARS-CoV-2 clades recovered a slight but significant decrease in estimated transmissibility over time for 10 of 13 clades analysed, largely corresponding to the accumulation of mutations associated with lower CEGA scores. Such an observation is consistent with the expectation of a decay in transmissibility for non-recombining lineages caused by the accumulation of deleterious mutations (often referred to as ‘Muller’s ratchet’)^57,58^. Given the high prevalence of C→T homoplasies, one plausible contributing factor is the accumulation of mutations over time due to host-editing activity, with deamination of cytidines being a hallmark of the APOBEC family of proteins^81^. However, such a small and gradual decay in transmissibility of existing lineages provides only part of the story. Lineage replacement dynamics are clearly driven by a combination of other phenomena, including the ‘sudden’ emergence of more transmissible lineages (such as Alpha and Delta) and the impact of pharmaceutical and non-pharmaceutical interventions on certain geographically restricted lineages.

Our analyses take advantage of the unprecedented size of the GISAID dataset^5,6^ thanks to the efforts of large numbers of contributing laboratories generating and openly sharing data, and aided by release of global phylogenies via the GISAID Audacity platform. While the data should be fairly representative of the extant diversity of SARS-CoV-2 in circulation globally, it is still affected by significant sampling heterogeneity. This arises primarily through variable sequencing efforts by different countries, and to a lesser extent from targeted sequencing of VoIs and VoCs after their identification. However, we reason that our approach should be largely robust to such sampling heterogeneities. Indeed, we rely on multiple replicates provided by the independent emergences of mutations and deletions in different genetic backgrounds. The fact that our results remain largely unaffected whether we compute our transmissibility scores using the mean or the median of the CEGA values obtained from all testable independent emergences of each mutation or deletion suggests our method produces well-behaved scores.

Recurrent mutations may also be detected erroneously in poorly resolved phylogenies. Uncertainty in the topology of SARS-CoV-2 phylogeny is difficult to assess and quantify because of the limited available genetic diversity which implies that most of the nodes of the phylogeny are supported by a sole SNV. Despite these characteristics, deep nodes of the phylogeny appear stable across trees produced by many different methods^11^. The use of parsimony to place new sequences into existing trees^63^, which is used by the GISAID Audacity platform, is estimated to place 97.2% of SARS-CoV-2 samples correctly with the initial placement, and the Audacity pipeline further optimises these placements using pseudo-Maximum Likelihood implemented in FastTree2^82^. To bias the CEGA score of any given mutation, errors in the topology would have to cause strains bearing the mutation in question to be systematically misplaced by groups of at least five (one of our filtering criteria) in lineages having a sister lineage not carrying the mutation. Given these parameters, our analysis should be robust to most phylogenetic uncertainty, because such uncertainty would tend to mix samples from related lineages and cause them to fail our pre-selection filters. Low levels of recombination may also result in the detection of recurrent mutations. However, such cases would still provide an assessment of the effect of that mutation in the recombinant lineage background, with our method largely agnostic to the mechanisms underlying how a recurrent mutation arises.

Transmissibility is a complex phenotypic trait and we have only analysed one aspect, namely the contribution from individual recurrent mutations and deletions. Although we expect our scores to be relatively robust to many sources of bias, it is important to stress that the transmissibility estimates for individual mutations and deletions, as well as for whole genomes, represent relative rather than absolute estimates (i.e., ‘additional’ changes in transmission). The values are also affected by choices during the normalisation procedure, including the value selected for generation time. They cannot therefore be simply compared to transmissibility estimates derived from other approaches. In addition, the consistent trend we recover of a slight decrease in poly-CEGA scores over time could conceivably arise from a systematic bias; although we have not been able to identify any plausible cause.

In summary, we herein make use of an extensive genomic dataset of SARS-CoV-2 to assess the contributions of recurrent mutations and deletions to the estimated transmissibility of SARS-CoV-2 through time. The per-generation scoring metric we developed highlights a transmissibility increase associated with mutations and deletions including those in the spike protein that were previously known or suspected to affect transmissibility, but also sheds light on the potential role of a wider set of mutations in other proteins. More fundamentally, generation of genome-wide estimates of transmissibility based on the individual contributions of mutations and deletions allows recovery of marked selective shifts in the transmissibility of SARS-CoV-2. In addition, we identify a tendency for phylogenetic clades to slightly decline in transmissibility through time after their emergence through the accumulation of mutations estimated to be weakly deleterious with respect to transmissibility. Such a trend may, in part, contribute to the patterns of lineage dynamics observed over the COVID-19 pandemic thus far.

## Methods

### Data acquisition

We downloaded the SARS-CoV-2 phylogeny provided by GISAID^5,6^ to registered users for data current to 29/08/2021 via the Audacity feature which optimises trees with a process based on the pipeline presented at https://github.com/roblanf/sarscov2phylo. The GISAID Audacity workflow aligns high coverage sequences to hCoV-19/Wuhan/WIV04/2019 (WIV04) following masking of all problematic sites^11^, as listed at https://github.com/W-L/ProblematicSites_SARS-CoV2 (accessed 29/08/2021). From the total alignment 400,000 distinct sequences are sampled taking 90% of the most recently submitted sequences, and a random selection to make up the rest. We additionally restricted our study to the assemblies available in the Mafft-based multiple sequence alignment available on the GISAID EpiCoV™ database (https://www.gisaid.org) already filtered to exclude samples displaying a total genome length <= 29,000bp, a fraction of ‘N’ nucleotides >5%, or displaying long series of leading or trailing ‘N’ nucleotides. We also discarded strains displaying spurious alleles at positions of known nucleotide deletions: Δ686-694; Δ11,288-11,296; Δ21,765-21,770; Δ21,991-21,993; Δ22,029-22,034; Δ28,248-28,253 (corresponding to NSP1 Δ141-143; NSP6 Δ106-108; S Δ69-70; S Δ144; S Δ156-157 and ORF8 Δ119-120 respectively). All ambiguous sites in the alignment were set to ‘N’. This resulted in an alignment and a maximum likelihood phylogeny both comprising 491,449 SARS-CoV-2 assemblies for downstream analysis. A full metadata table, acknowledgement of data generators and submitters and accession exclusions is provided in Supplementary Table S5.

### Detection of recurrent mutational events

We filtered the alignment to only include variable positions for which the most common alternate allele was present in at least 0.1% of accessions. The heterogeneous quality of assemblies submitted to GISAID poses challenges to automated calling of deletions. We therefore restricted our study to the six deletions listed above and checked them manually in the alignment. We additionally masked sequencing error prone and/or back-mutation prone nucleotide positions 11,083 and 21,575^11^ as well as the first 150 and last 300 bp because of their higher proportion of missing data. We applied HomoplasyFinder v0.0.0.9^83^ to this alignment using the GISAID Audacity tree to quantify the number of independent emergences of all mutations and deletions considered and to identify the parental node of every recurrent mutation/deletion in the dataset. This resulted in detection of 1,552 total homoplastic positions. We discarded all homoplasies for which we did not have at least five independent emergences supported by nodes obeying the following rules: (i) at least ten descending tips carrying either allele, (ii) no children node passing (i) embedded carrying a subsequent mutation at the same site and (iii) each descending lineage of the considered node must show an allele frequency >90%. This latter rule prevents taking into account of spurious clades with series of back-and-forth mutations at the same position. We previously showed that such filtering was necessary to obtain robust and symmetrically distributed scores^9^. The homoplasy detection and filtration procedure has been thoroughly described in our first implementation of a Ratio of Homoplastic Offspring (RoHO) scoring method (see Methods of van Dorp and Richard et al 2020^9^). Following each of these steps 719 homoplasies passed the filtering criterion. While this does not include all mutations flagged as of concern, for instance in known VoCs and VoIs, further mutations may pass our stringency filters as additional genomes are sequenced and shared.

### Computation of the Coefficients of Exponential Growth Increase (CEGAs)

Our previous work^9^ considered the ratio of homoplastic offspring (‘RoHO’) of two lineages descending from a given ancestor (the number of genomes carrying and not carrying the considered mutation). While RoHO can capture accurately the increased success of a lineage compared to its sister lineage, it fails to account for the fact that under an epidemic growth model, RoHO is not expected to be constant over time. Here we outline a metric that accounts for this variation in time, which we call the Coefficients of Exponential Growth Alteration (CEGAs). This approach aims to normalise the excess of the number of offspring per generation under exponential growth (Supplementary Figure S5).

We first consider the general case of two lineages, a wildtype (w) and a variant (v). If the wildtype lineage has a mean generation interval 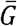 with a coefficient of variation *κ* then the reproduction number *R* is related to the growth rate *r* by

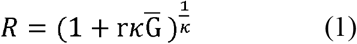

The population size of the wildtype after time *t* is given by

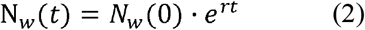

If we assume that the variant has an increased reproduction number *R*_*v*_ = *ρR*, this means that

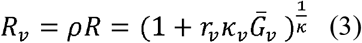

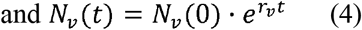

For the usual case of comparing two lineages (e.g. Davies *et al*. 2021^84^, Park *et al*. 2021^85^), the instantaneous proportion of isolates from the variant over time *p*(t) = *N*_*v*_/(*N*_*v+*_ *N*_*w*_) provides a means to estimate the relative transmissibility of the variant, under the assumption that sampling is random.

Here, we wish to calculate the effect of a particular recurrent mutation or deletion *V* on transmissibility. *V* may have emerged on multiple occasions, each representing an opportunity to estimate this effect. Consider a particular homoplastic node in the phylogeny where *V* has emerged, producing a clade comparison between two lineages: a wildtype without *V* and a variant lineage with *V*. We make the simplifying assumption that the generation interval is fixed (*k*_*v*_ = *k = 0*) and that *V* has no effect on the generation interval 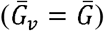. In this case, the equations relating reproduction number, growth rate, and generation interval for the two lineages at a given clade comparison simplify to:

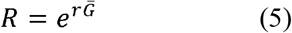

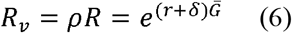

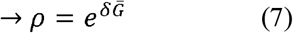

We assume that *V* confers a growth rate advantage of *r*_*v*_ = *r + δ* compared to the basal growth rate *r* This means that *δ* is the additive growth rate due to *V* for a given clade comparison, with units of per time. Theoretically both *r* and *δ* can differ between clade comparisons, due for example to different effects of *V* in different genetic backgrounds (epistasis) or differences in epidemiological setting (which could cause differences in generation interval). When *δ* > 0, the variant lineage produces a larger number of offspring so its population will grow faster with time (and vice versa for. *δ* < 0). From the phylogeny, we can measure the proportion of tips that are from the variant lineage in this clade comparison. The problem is that this simple proportion is not the instantaneous proportion *p*(*t*) but the cumulative proportion P(T). We must relate this phylogenetic information to the population dynamics to estimate *δ*.

Our previous ‘RoHO’ index^9^ (the ratio of tips in each lineage) does not normalise by time. Assuming random sampling, the proportion of tips in each lineage after a time T will be equivalent to the cumulative proportion of the populations P(T), given by

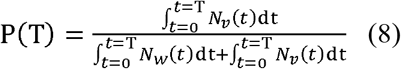

Substituting in equations (2) and (4) and simplifying, we obtain

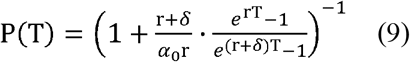

Here we have defined the initial population ratio 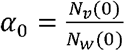. Because we are dealing with a split in the phylogeny from a single ancestral node, we are assuming that both lineages start with an equal population size and thus have *α*_*0*_ = 1. Note that equation (9) has dependence on r, since in general the cumulative proportion of tips comparing two lineages with/without *V* depends both on the basal growth rate (r)and the additional effect of *V*(*δ*).

However, typically the clade comparisons we are using involve very closely-related isolates which are by virtue ‘shallow’ in time. We can therefore approximate this expression by noting that the Taylor expansion for small T is

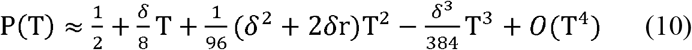

If we assume that *δ* ≪ 1 then under the additional condition rT ≪ 6 we can drop terms of O(T^2^) and higher, so equation (10) approximates to

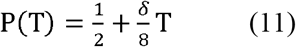

We can now rearrange this expression for*δ*

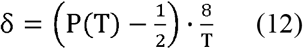

We estimate T as 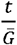 where *t* is the timespan between the oldest and the youngest offspring of the considered node. We use 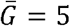 days based on the mean and median estimates provided by four independent studies^86–89^. Values of *δ* (‘CEGA scores’) presented in the manuscript are therefore given per generation interval. For a given mutation or deletion *V* we obtain multiple CEGA scores from different clade comparisons in the phylogeny. We summarise these using the mean and median of these CEGA scores (provided in Supplementary Table S3). Although this method allows for *δ* to differ between clade comparisons, we average *δ* over these comparisons to get an average *δ* for each *V* and interpret this as the expected average effect of *V* on growth rate.

In summary, equation (12) allows for a rough quantitative comparison of the transmissibility effects of different recurrent mutations or deletions in a relative sense, using the information available in a phylogeny while accounting for variation in time. One alternative approach would be to try to estimate the instantaneous proportion with a sliding time window over the phylogeny, but the short timescales involved and the unreliability of sample dates means that we do not think this is feasible at this stage. We stress that we have made several simplifying assumptions to arrive at this expression.

### Estimation of the transmissibility of major SARS-CoV-2 clades

Our method estimates a CEGA value for each recurrent mutation and deletion under scrutiny (passing filters) in the alignment. It follows that the estimated transmissibility of any individual genome is influenced by the multiple mutations or deletions it carries. To test a multilocus implementation of the model we estimated the transmissibility score for each genome in our dataset which we term poly-CEGA. To do so we employed a simple model, where the effect of mutations and deletions on transmissibility are considered as independent, by computing the sum of mean CEGA scores of a genome made up of mutations 1…*n* as poly-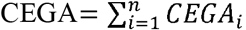, where sites for which no CEGA score was computed (did not pass our filters) are considered as having a CEGA of 0. By extension, we assessed the combined transmissibility change estimate – the poly-CEGA – of all SARS-CoV-2 isolates assessed over the global population and by NextStrain clades for those including > 1,000 genomes.

## Acknowledgements

This analysis was enabled by the GISAID EpiCoV™ database (https://www.gisaid.org) and regular Audacity releases. We gratefully acknowledge all contributing and submitting laboratories around the globe including the COG-UK consortium (https://www.cogconsortium.uk) who have openly shared large numbers of UK SARS-CoV-2 assemblies. A full list of acknowledgements of submitting and originating laboratories is provided in Supplementary Table S5. We are also grateful to developers and maintainers of CoVariants (https://covariants.org) and Outbreak.info (https://outbreak.info) for information on mutation and deletion carriage in VoCs and VoIs. We additionally thank Sang Woo Park for helpful correspondence and discussion of the derivation of the CEGA metric.

## References

1. van Dorp, L. et al. Emergence of genomic diversity and recurrent mutations in SARS-CoV-2. Infect. Genet. E vol. 83, 104351 (2020).

2. Kumar, S. et al. An evolutionary portrait of the progenitor SARS-CoV-2 and its dominant offshoots in COVID-19 pandemic. Mol. Biol. Evol. 38, 3046–3059 (2021).

3. Rambaut, A. Phylogenetic analysis of 23 nCoV-2019 genomes, 2020-01-23 - SARS-CoV-2 coronavirus / nCoV-2019 Genomic Epidemiology - Virological. Virological https://virological.org/t/phylogenetic-analysis-of-23-ncov-2019-genomes-2020-01-23/335 (2020).

4. Hadfield, J. et al. Nextstrain: real-time tracking of pathogen evolution. Bioinformatics 34, 4121–4123 (2018).

5. Shu, Y. & McCauley, J. GISAID: Global initiative on sharing all influenza data – from vision to reality. Eurosurveillance 22, 30494 (2017).

6. Elbe, S. & Buckland-Merrett, G. Data, disease and diplomacy: GISAID’s innovative contribution to global health. Glob. Challenges 1, 33–46 (2017).

7. Singer, J., Gifford, R., Cotten, M. & Robertson, D. CoV-GLUE: A Web Application for Tracking SARS-CoV-2 Genomic Variation. (2020) doi:10.20944/PREPRINTS202006.0225.V1.

8. McCarthy, K. R. et al. Recurrent deletions in the SARS-CoV-2 spike glycoprotein drive antibody escape. Science (80-.). 371, 1139–1142 (2021).

9. van Dorp, L. et al. No evidence for increased transmissibility from recurrent mutations in SARS-CoV-2. Nat. Commun. 11, 5986 (2020).

10. Kemp, S. et al. Recurrent emergence and transmission of a SARS-CoV-2 Spike deletion ΔH69/V70. Cell Rep. 35, 109292 (2021).

11. Turakhia, Y. et al. Stability of SARS-CoV-2 phylogenies. PLOS Genet. 16, e1009175 (2020).

12. Martin, D. P. et al. The emergence and ongoing convergent evolution of the N501Y lineages coincides with a major global shift in the SARS-CoV-2 selective landscape. Cell (2021) doi:10.1101/2021.02.23.21252268.

13. Duffy, S., Shackelton, L. A. & Holmes, E. C. Rates of evolutionary change in viruses: patterns and determinants. Nat. Rev. Genet. 9, 267–276 (2008).

14. Simmonds, P. Rampant C→U Hypermutation in the Genomes of SARS-CoV-2 and Other Coronaviruses: Causes and Consequences for Their Short-and Long-Term Evolutionary Trajectories. mSphere 5, (2020).

15. Giorgio, S. Di, Martignano, F., Torcia, M. G., Mattiuz, G. & Conticello, S. G. Evidence for host-dependent RNA editing in the transcriptome of SARS-CoV-2. Sci. Adv. 6, eabb5813 (2020).

16. De Maio, N. et al. Mutation rates and selection on synonymous mutations in SARS-CoV-2. Genome Biol. Evol. 13, 2021.01.14.426705 (2021).

17. Mourier, T. et al. Host-directed editing of the SARS-CoV-2 genome. Biochem. Biophys. Res. Commun. (2020) doi:10.1016/j.bbrc.2020.10.092.

18. Lythgoe, K. A. et al. SARS-CoV-2 within-host diversity and transmission. Science (2021) doi:10.1126/science.abg0821.

19. Richard, D., Owen, C. J., Dorp, L. van & Balloux, F. No detectable signal for ongoing genetic recombination in SARS-CoV-2. bioRxiv 2020.12.15.422866 (2020) doi:10.1101/2020.12.15.422866.

20. VanInsberghe, D., Neish, A. S., Lowen, A. C. & Koelle, K. Recombinant SARS-CoV-2 genomes are currently circulating at low levels. bioRxiv (2021).

21. Varabyou, A., Pockrandt, C., Salzberg, S. L. & Pertea, M. Rapid detection of inter-clade recombination in SARS-CoV-2 with Bolotie. bioRxiv 2020.09.21.300913 (2020) doi:10.1101/2020.09.21.300913.

22. Jackson, B. et al. Recombinant SARS-CoV-2 genomes involving lineage B.1.1.7 in the UK - SARS-CoV-2 coronavirus / SARS-CoV-2 Molecular Evolution - Virological. Virological https://virological.org/t/recombinant-sars-cov-2-genomes-involving-lineage-b-1-1-7-in-the-uk/658 (2021).

23. Jackson, B. et al. Generation and transmission of interlineage recombinants in the SARS-CoV-2 pandemic. Cell (2021) doi:10.1016/j.cell.2021.08.014.

24. Turkahia, Y. et al. Pandemic-Scale Phylogenomics Reveals Elevated Recombination Rates in the SARS-CoV-2 Spike Region. bioRxiv 2021.08.04.455157 (2021) doi:10.1101/2021.08.04.455157.

25. McCallum, M. et al. N-terminal domain antigenic mapping reveals a site of vulnerability for SARS-CoV-2. Cell 184, 2332–2347.e16 (2021).

26. Andreano, E. et al. SARS-CoV-2 escape in vitro from a highly neutralizing COVID-19 convalescent plasma. bioRxiv 2020.12.28.424451 (2020) doi:10.1101/2020.12.28.424451.

27. Korber, B. et al. Tracking Changes in SARS-CoV-2 Spike: Evidence that D614G Increases Infectivity of the COVID-19 Virus. Cell 182, 812–827.e19 (2020).

28. Volz, E. et al. Evaluating the Effects of SARS-CoV-2 Spike Mutation D614G on Transmissibility and Pathogenicity. Cell 184, 64–75.e11 (2021).

29. van Dorp, L., Houldcroft, C. J., Richard, D. & Balloux, F. COVID-19, the first pandemic in the post-genomic era. Current Opinion in Virology vol. 50 40–48 (2021).

30. Rambaut, A. et al. Preliminary genomic characterisation of an emergent SARS-CoV-2 lineage in the UK defined by a novel set of spike mutations. https://virological.org/t/preliminary-genomic-characterisation-of-an-emergent-sars-cov-2-lineage-in-the-uk-defined-by-a-novel-set-of-spike-mutations/563 (2020).

31. Tegally, H. et al. Detection of a SARS-CoV-2 variant of concern in South Africa. Nature 592, 438–443 (2021).

32. Faria, N. R. et al. Genomics and epidemiology of the P.1 SARS-CoV-2 lineage in Manaus, Brazil. Science. eabh2644 (2021) doi:10.1126/science.abh2644.

33. Mlcochova, P. et al. SARS-CoV-2 B.1.617.2 Delta variant emergence and vaccine breakthrough. bioRxiv 2021.05.08.443253 (2021).

34. Piccoli, L. et al. Mapping Neutralizing and Immunodominant Sites on the SARS-CoV-2 Spike Receptor-Binding Domain by Structure-Guided High-Resolution Serology. Cell 183, 1024–1042.e21 (2020).

35. Harvey, W. T. et al. SARS-CoV-2 variants, spike mutations and immune escape. Nature Reviews Microbiology vol. 19 409–424 (2021).

36. Kemp, S. A. et al. Neutralising antibodies drive Spike mediated SARS-CoV-2 evasion. Nature 592, 277–282 (2021).

37. Avanzato, V. A. et al. Case Study: Prolonged Infectious SARS-CoV-2 Shedding from an Asymptomatic Immunocompromised Individual with Cancer. Cell 183, 1901–1912.e9 (2020).

38. Choi, B. et al. Persistence and Evolution of SARS-CoV-2 in an Immunocompromised Host. N. Engl. J. Med. 383, (2020).

39. Starr, T. N. et al. Deep Mutational Scanning of SARS-CoV-2 Receptor Binding Domain Reveals Constraints on Folding and ACE2 Binding. Cell 182, 1295–1310.e20 (2020).

40. Zahradník, J. et al. SARS-CoV-2 variant prediction and antiviral drug design are enabled by RBD in vitro evolution. Nat. Microbiol. 6, 1188–1198 (2021).

41. Nelson, G. et al. Molecular dynamic simulation reveals E484K mutation enhances spike RBD-ACE2 affinity and the combination of E484K, K417N and N501Y mutations (501Y.V2 variant) induces conformational change greater than N501Y mutant alone, potentially resulting in an escap. bioRxiv 2021.01.13.426558 (2021) doi:10.1101/2021.01.13.426558.

42. Volz, E. et al. Transmission of SARS-CoV-2 Lineage B.1.1.7 in England: Insights from linking epidemiological and genetic data. medRxiv 2020.12.30.20249034 (2021) doi:10.1101/2020.12.30.20249034.

43. Li, B. et al. Viral infection and transmission in a large, well-traced outbreak caused by the SARS-CoV-2 Delta variant. medRxiv 2021.07.07.21260122 (2021) doi:10.1101/2021.07.07.21260122.

44. Garcia-Beltran, W. F. et al. Multiple SARS-CoV-2 variants escape neutralization by vaccine-induced humoral immunity. Cell 184, 2372–2383 (2021).

45. Cele, S. et al. Escape of SARS-CoV-2 501Y.V2 from neutralization by convalescent plasma. Nature 593, 142–146 (2021).

46. Wibmer, C. K. et al. SARS-CoV-2 501Y.V2 escapes neutralization by South African COVID-19 donor plasma. Nat. Med. 1–4 (2021) doi:10.1038/s41591-021-01285-x.

47. Wu, K. et al. mRNA-1273 vaccine induces neutralizing antibodies against spike mutants from global SARS-CoV-2 variants. bioRxiv Prepr. Serv. Biol. 2021.01.25.427948 (2021) doi:10.1101/2021.01.25.427948.

48. Hoffmann, M. et al. SARS-CoV-2 variants B.1.351 and B.1.1.248: Escape from therapeutic antibodies and antibodies induced by infection and vaccination. Cell 184, 2384–2393 (2021).

49. Bernal, J. L. et al. Effectiveness of COVID-19 vaccines against the B.1.617.2 variant. New Engl. J. Med. 385, 585–594 (2021).

50. Bager, P. et al. Increased Risk of Hospitalisation Associated with Infection with SARS-CoV-2 Lineage B.1.1.7 in Denmark. SSRN Electron. J. (2021) doi:10.2139/ssrn.3792894.

51. Davies, N. G. et al. Increased mortality in community-tested cases of SARS-CoV-2 lineage B.1.1.7. Nature 1–5 (2021) doi:10.1038/s41586-021-03426-1.

52. Graham, M. S. et al. Changes in symptomatology, reinfection, and transmissibility associated with the SARS-CoV-2 variant B.1.1.7: an ecological study. Lancet Public Heal. (2021) doi:10.1016/s2468-2667(21)00055-4.

53. Santos De Oliveira, M. H., Lippi, G. & Henry, B. M. Sudden rise in COVID-19 case fatality among young and middle-aged adults in the south of Brazil after identification of the novel B.1.1.28.1 (P.1) SARS-CoV-2 strain: analysis of data from the state of Parana. medRxiv 2021.03.24.21254046 (2021) doi:10.1101/2021.03.24.21254046.

54. Bager, P., Wohlfahrt, J., Rasmussen, M., Albertsen, M. & Krause, T. G. Hospitalisation associated with SARS-CoV-2 delta variant in Denmark. Lancet Infect. Dis. (2021) doi:10.1016/S1473-3099(21)00580-6.

55. Twohig, K. A. et al. Hospital admission and emergency care attendance risk for SARS-CoV-2 delta (B.1.617.2) compared with alpha (B.1.1.7) variants of concern: a cohort study. Lancet Infect. Dis. (2021) doi:10.1016/S1473-3099(21)00475-8.

56. Stern, A. et al. The unique evolutionary dynamics of the SARS-CoV-2 Delta variant Israel Consortium of SARS-CoV-2 sequencing. medRxiv 2021.08.05.21261642 (2021) doi:10.1101/2021.08.05.21261642.

57. Felsenstein, J. The evolution advantage of recombination. Genetics 78, 737–756 (1974).

58. Muller, H. J. The relation of recombination to mutational advance. Mutat. Res. - Fundam. Mol. Mech. Mutagen. 1, 2–9 (1964).

59. du Plessis, L. et al. Establishment and lineage dynamics of the SARS-CoV-2 epidemic in the UK. Science (80-.). 10, eabf2946 (2021).

60. Duchêne, S. & Lanfear, R. Phylogenetic uncertainty can bias the number of evolutionary transitions estimated from ancestral state reconstruction methods. J. Exp. Zool. Part B Mol. Dev. E vol. 324, 517–524 (2015).

61. Thomson, E. C. et al. Circulating SARS-CoV-2 spike N439K variants maintain fitness while evading antibody-mediated immunity. Cell 184, 1171–1187.e20 (2021).

62. Hodcroft, E. B. et al. Spread of a SARS-CoV-2 variant through Europe in the summer of 2020. Nature vol. 595 707–712 (2021).

63. Yatish Turakhia, et al. Ultrafast Sample Placement on Existing Trees (UShER) Empowers Real-Time Phylogenetics for the SARS-CoV-2 Pandemic. Nat. Genet. 1348, (2021).

64. Greaney, A. J. et al. Comprehensive mapping of mutations in the SARS-CoV-2 receptor-binding domain that affect recognition by polyclonal human plasma antibodies. Cell Host Microbe 29, 463–476.e6 (2021).

65. Nonaka, C. K. V. et al. Genomic Evidence of SARS-CoV-2 Reinfection Involving E484K Spike Mutation, Brazil. Emerg. Infect. Dis. 27, (2021).

66. Alaa Abdel Latif et al. N:D3L Mutation Report http://Outbreak.info. (2021).

67. Matthew Parker, A. D. et al. Altered Subgenomic RNA Expression in SARS-CoV-2 B.1.1.7. bioRxiv 2021.03.02.433156 (2021) doi:10.1101/2021.03.02.433156.

68. Johnson, B. A. et al. Furin Cleavage Site Is Key to SARS-CoV-2 Pathogenesis. bioRxiv Prepr. Serv. Biol. (2020) doi:10.1101/2020.08.26.268854.

69. Liu, Y. et al. Delta spike P681R mutation enhances SARS-CoV-2 fitness over Alpha variant 1. bioRxiv 2021.08.12.456173 (2021) doi:10.1101/2021.08.12.456173.

70. Gu, Z., Gu, L., Eils, R., Schlesner, M. & Brors, B. circlize implements and enhances circular visualization in R. Bioinformatics 30, 2811–2812 (2014).

71. Gel, B. & Serra, E. karyoploteR: an R/Bioconductor package to plot customizable genomes displaying arbitrary data. Bioinformatics 33, 3088–3090 (2017).

72. Lan, J. et al. Structure of the SARS-CoV-2 spike receptor-binding domain bound to the ACE2 receptor. Nature 581, 215–220 (2020).

73. Obermeyer, F. et al. Analysis of 2.1 million SARS-CoV-2 genomes identifies mutations associated with transmissibility. medRxiv 2021.09.07.21263228 (2021) doi:10.1101/2021.09.07.21263228.

74. Greaney, A. J. et al. Complete Mapping of Mutations to the SARS-CoV-2 Spike Receptor-Binding Domain that Escape Antibody Recognition. Cell Host Microbe 29, 44–57.e9 (2021).

75. Liu, Z. et al. Identification of SARS-CoV-2 spike mutations that attenuate monoclonal and serum antibody neutralization. (2021) doi:10.1016/j.chom.2021.01.014.

76. Zhang, Q. et al. Potent and protective IGHV3-53/3-66 public antibodies and their shared escape mutant on the spike of SARS-CoV-2. Nat. Commun. 12, (2021).

77. Zhou, D. et al. Evidence of escape of SARS-CoV-2 variant B.1.351 from natural and vaccine-induced sera. Cell 184, 2348–2361.e6 (2021).

78. Dicken, S. J. et al. Characterisation of B.1.1.7 and Pangolin coronavirus spike provides insights on the evolutionary trajectory of SARS-CoV-2. bioRxiv 2021.03.22.436468 (2021) doi:10.1101/2021.03.22.436468.

79. Eguia, R. T. et al. A human coronavirus evolves antigenically to escape antibody immunity. PLOS Pathog. 17, e1009453 (2021).

80. Kistler, K. E. & Bedford, T. Evidence for adaptive evolution in the receptor-binding domain of seasonal coronaviruses OC43 and 229E. Elife 10, 1–35 (2021).

81. Bishop, K. N., Holmes, R. K., Sheehy, A. M. & Malim, M. H. APOBEC-mediated editing of viral RNA. Science. 305, 645 (2004).

82. Price, M. N., Dehal, P. S. & Arkin, A. P. FastTree 2 – Approximately Maximum-Likelihood Trees for Large Alignments. PLoS One 5, e9490 (2010).

83. Crispell, J., Balaz, D. & Gordon, S. V. HomoplasyFinder: a simple tool to identify homoplasies on a phylogeny. Microb. genomics 5, (2019).

84. Davies, N. G. et al. Estimated transmissibility and impact of SARS-CoV-2 lineage B.1.1.7 in England. Science (80-.). 372, (2021).

85. Park, S. W. et al. Roles of generation-interval distributions in shaping relative epidemic strength, speed, and control of new SARS-CoV-2 variants. medRxiv 2021.05.03.21256545 (2021) doi:10.1101/2021.05.03.21256545.

86. Griffin, J. et al. Rapid review of available evidence on the serial interval and generation time of COVID-19. BMJ Open 10, 40263 (2020).

87. Ganyani, T. et al. Estimating the generation interval for coronavirus disease (COVID-19) based on symptom onset data, March 2020. Eurosurveillance 25, 1 (2020).

88. Ferretti, L. et al. Quantifying SARS-CoV-2 transmission suggests epidemic control with digital contact tracing. Science. 368, (2020).

89. Lehtinen, S., Ashcroft, P. & Bonhoeffer, S. On the relationship between serial interval, infectiousness profile and generation time. J. R. Soc. Interface 18, 20200756 (2021).

