## Supplementary material for "A phylogeny-based metric for estimating changes in transmissibility from recurrent mutations in SARS-CoV-2"

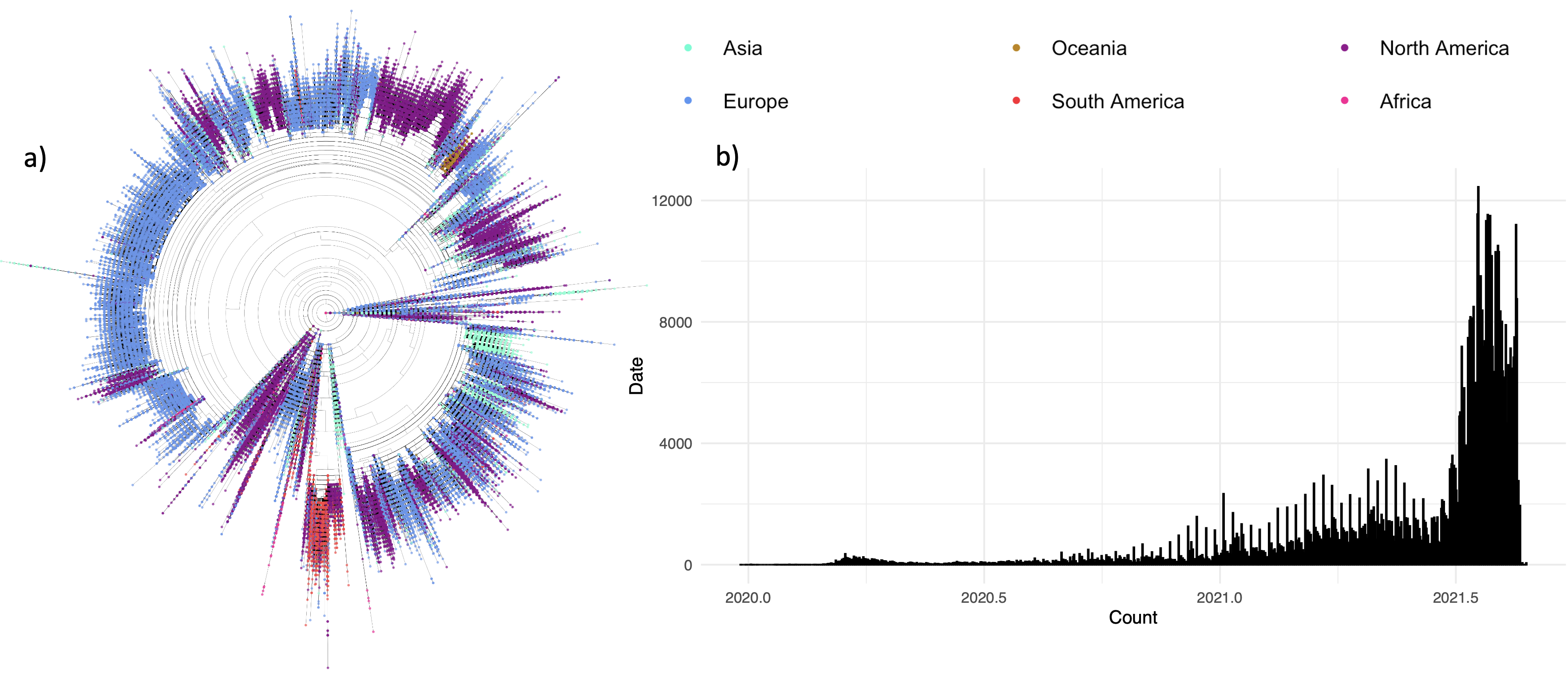

**Figure S1** (a) Maximum likelihood phylogenetic tree for 651,921 SARS-CoV-2 assemblies available via the GISAID Audacity pipeline, downloaded on 02/09/2021 with tips highlighted as per continental region as given in the legend (top right). (b) Temporal coverage of our dataset as seen by a histogram of the date of sample collection of all assemblies included in the dataset (using a bin size of 1000).

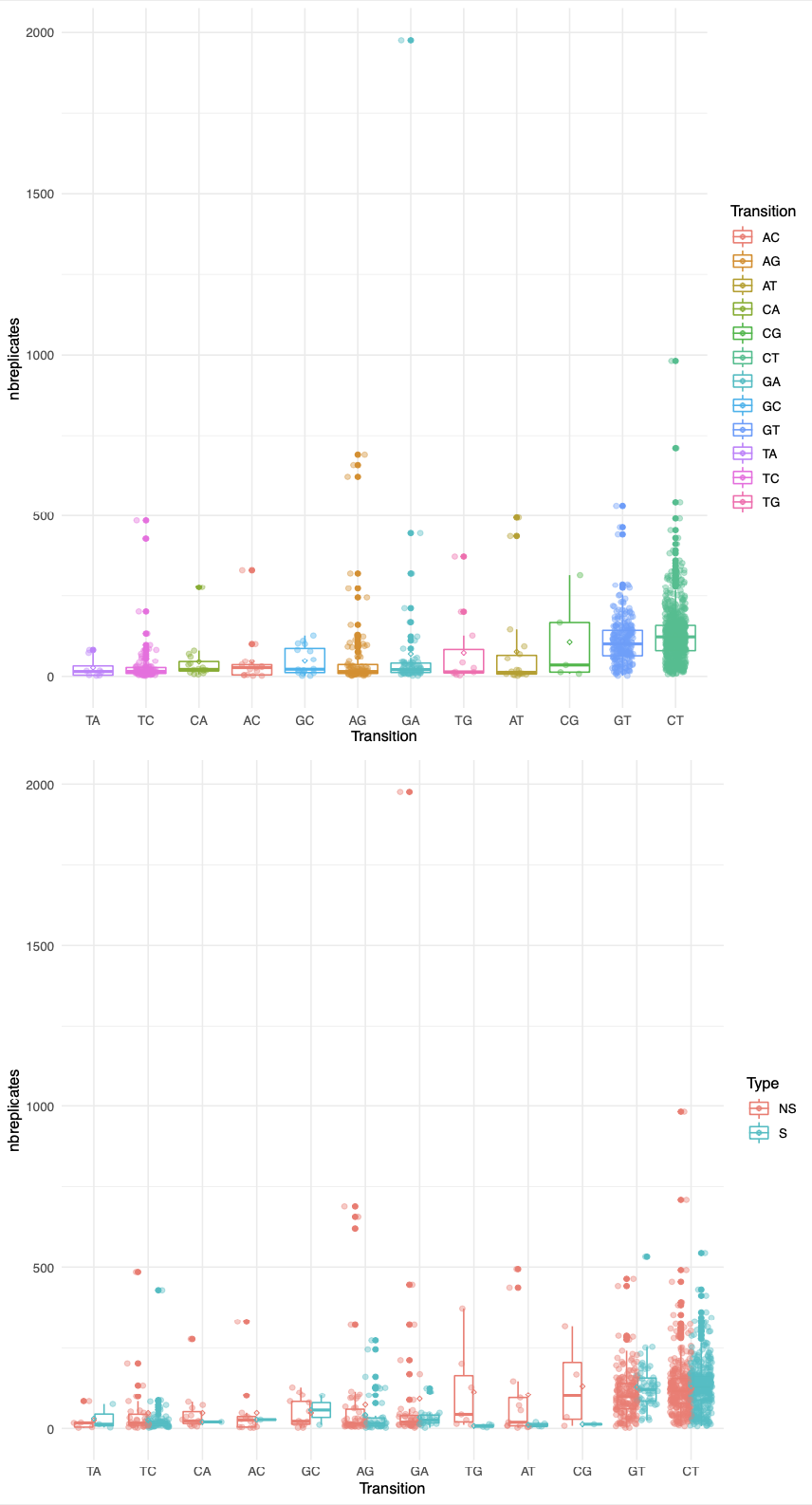

**Figure S2:** Boxplots providing the number of detected emergences (nbreplicates) for different transitions (CT equivalent to C🡪T etc.) observed in the SARS-CoV-2 alignment. The top plot provides the distributions at every site, with the lower providing the number of emergences estimated by position in the SARS-CoV-2 genome.

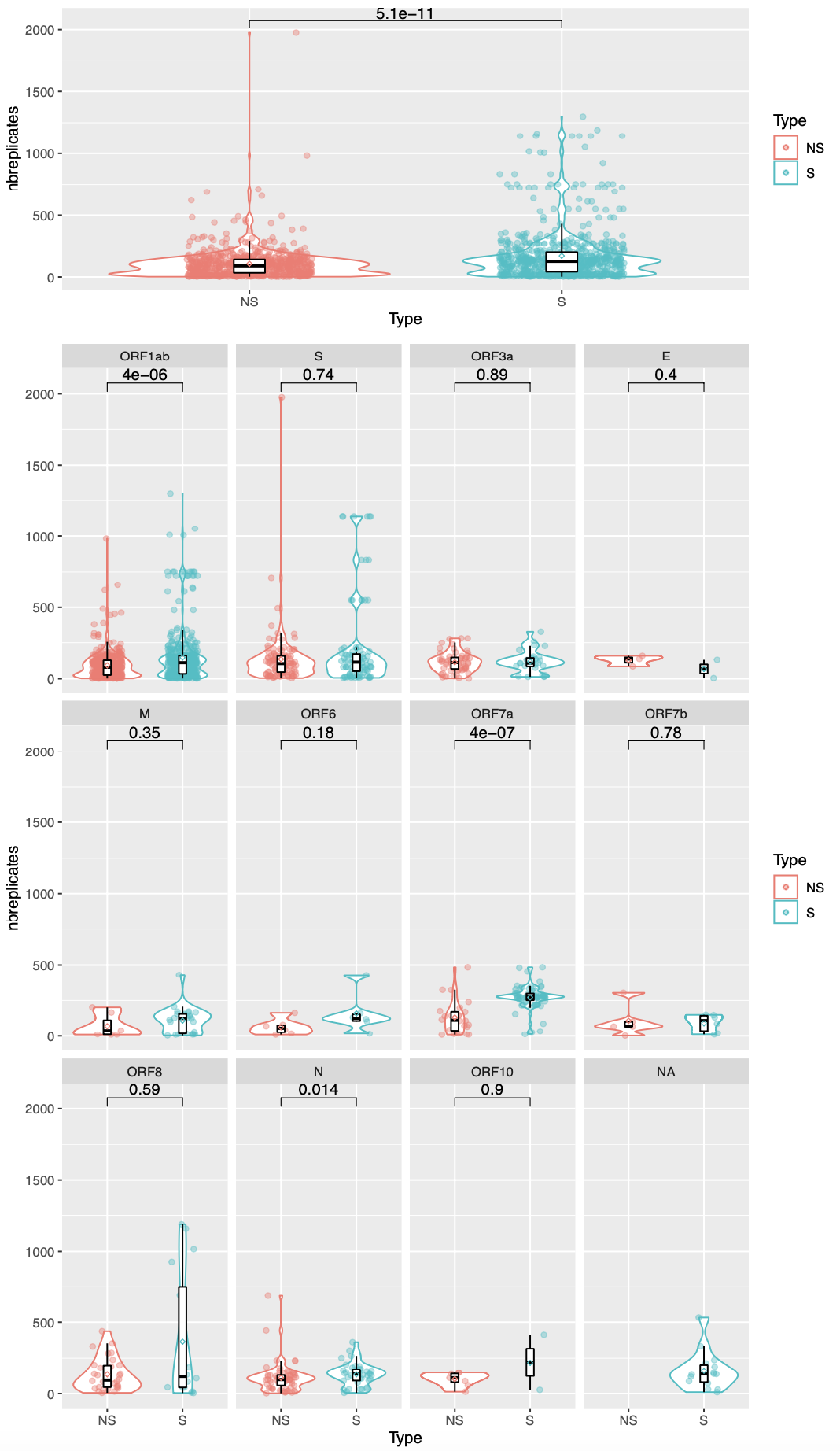

**Figure S3:** Boxplots providing the number of detected emergences (nbreplicates) associated to recurrent mutations at nonsynonymous (NS - orange) and synonymous (S - blue) sites. The top plot provides the distributions genome-wide, with the lower providing the number of emergences estimated by ORF in the SARS-CoV-2 genome. ‘NA’ refers to non-coding regions. *P*-values are provided following Wilcoxon test.

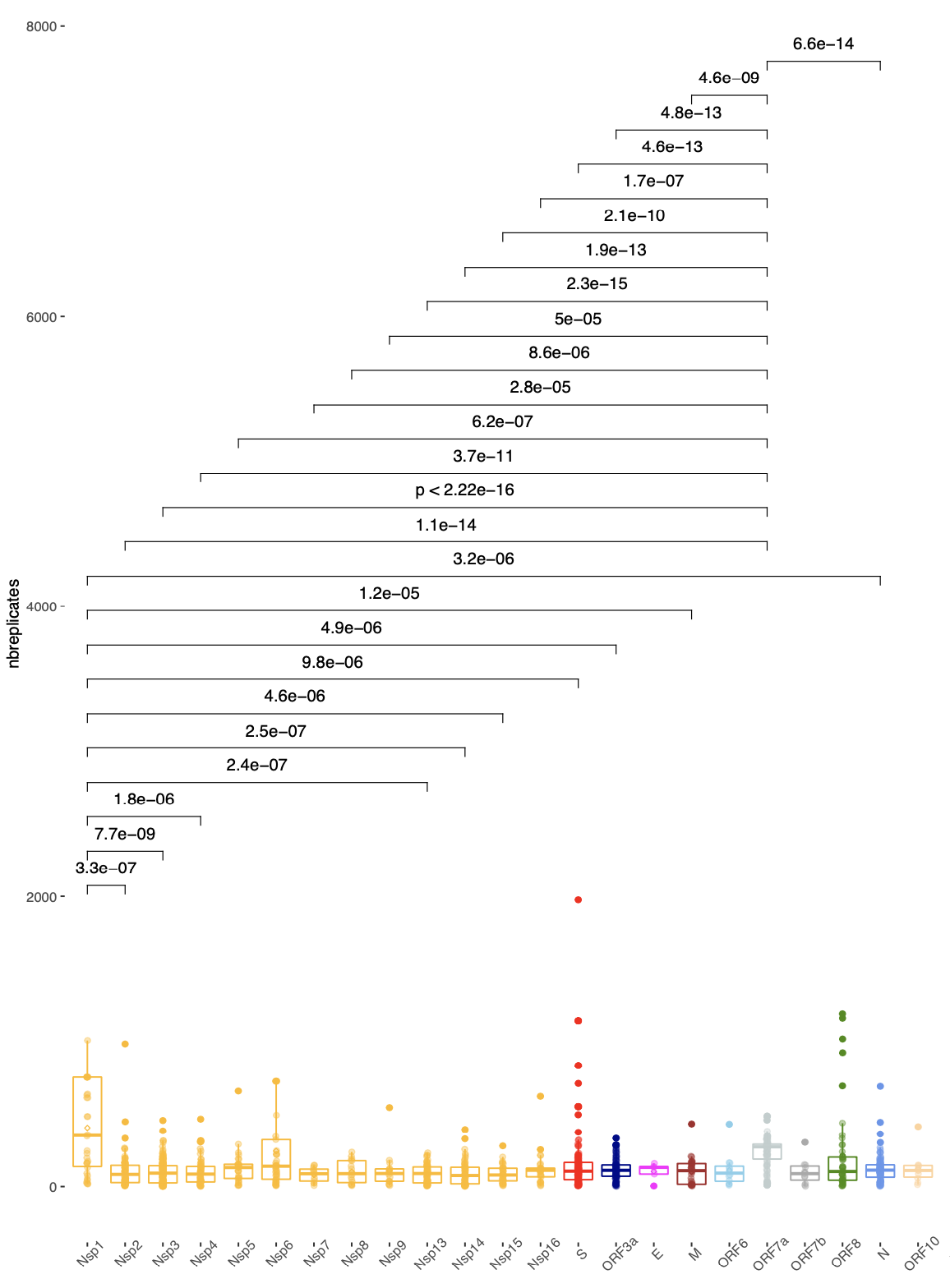

**Figure S4:** Boxplots providing the number of replicates (y-axis) associated to recurrent mutations at genomic features over the SARS-CoV-2 alignment. Pairwise comparisons between ORFs with a pairwise Wilcoxon *p*<0.0001 are demarcated.

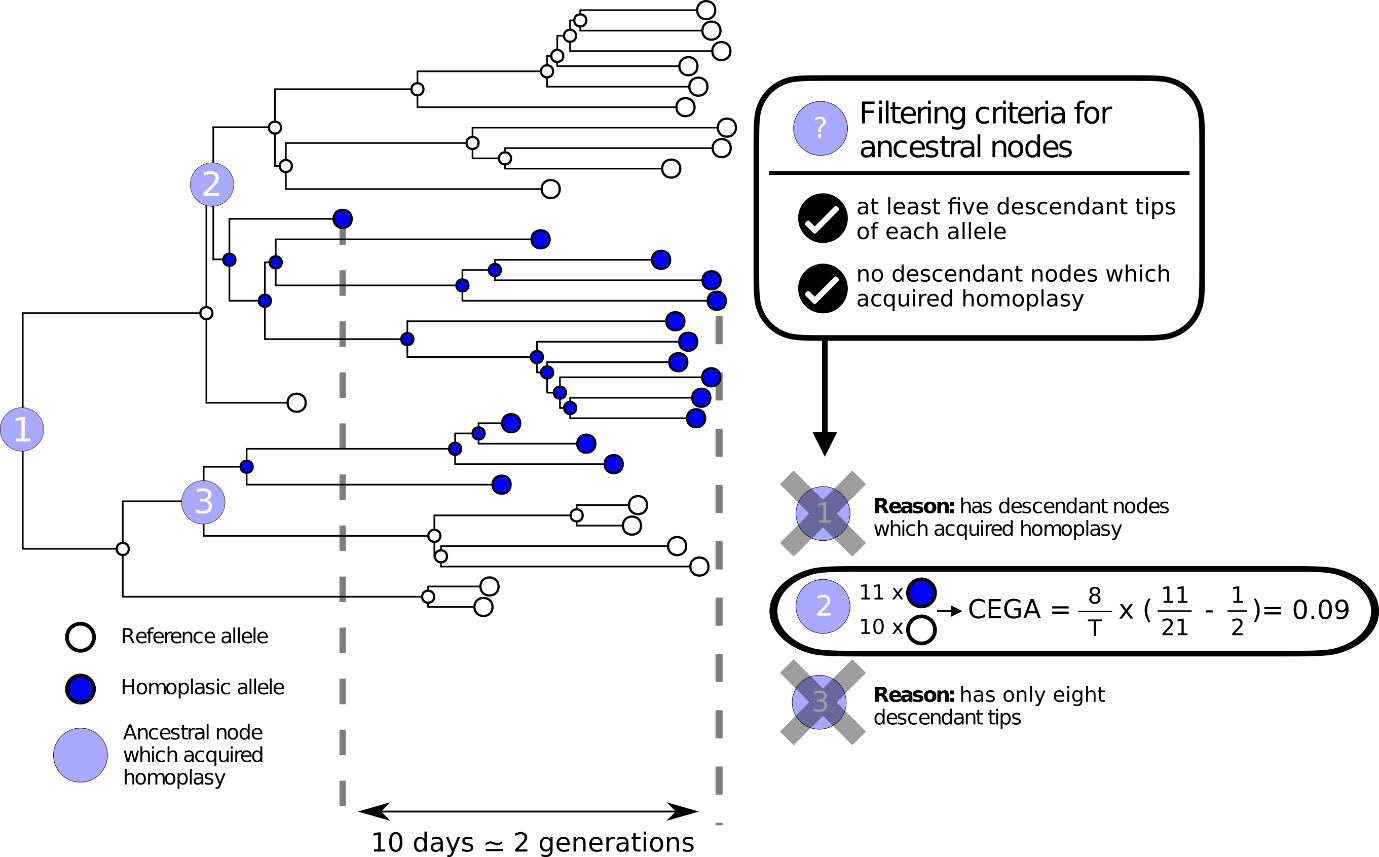

**Figure S5:** Schematic illustration of the rationale underlying the CEGA scoring metric.

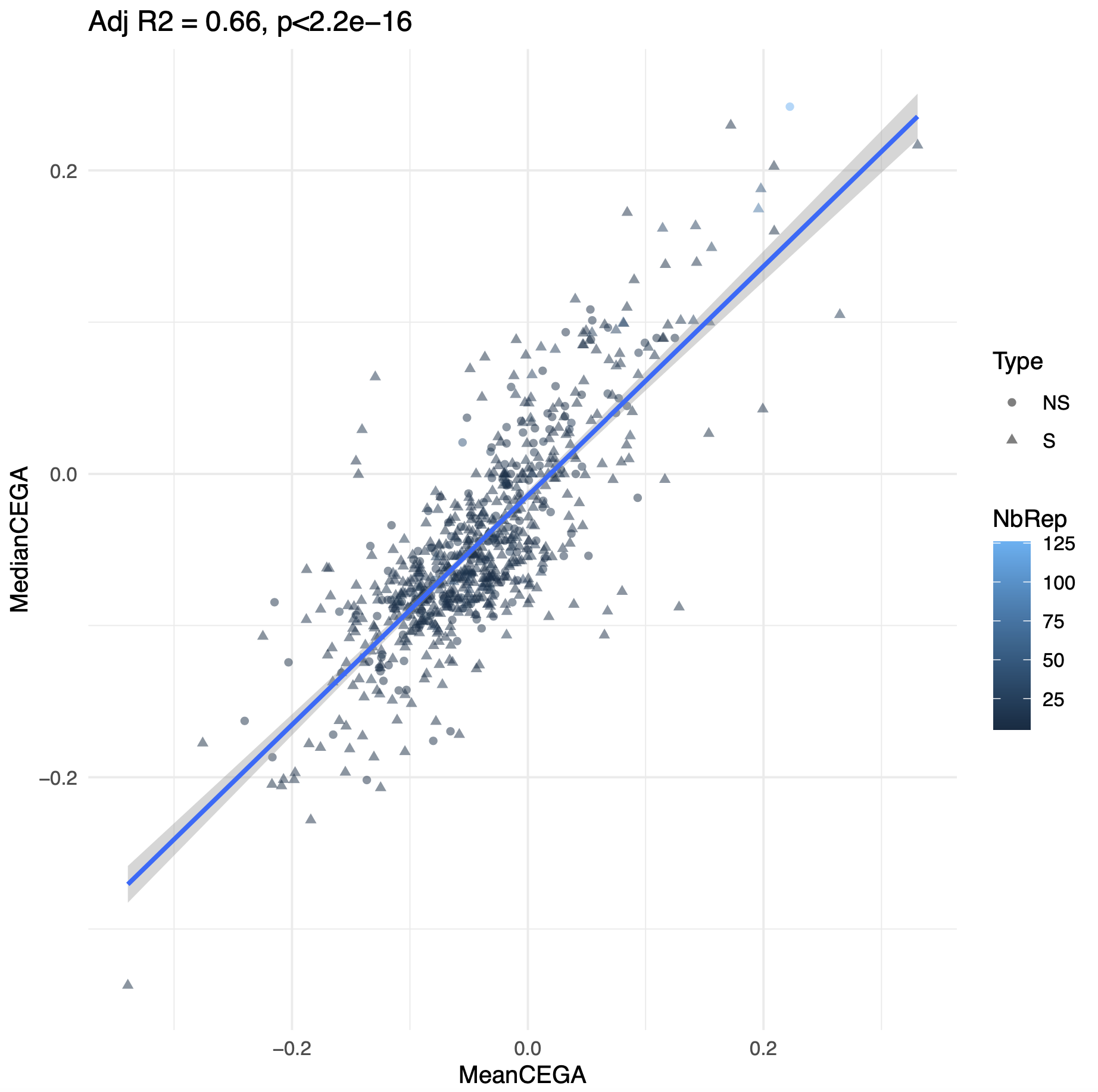

**Figure S6:** Concordance between mean and median CEGA values. Colour scale gives the number of associated replicates with point symbols denoting synonymous or nonsynonymous status.

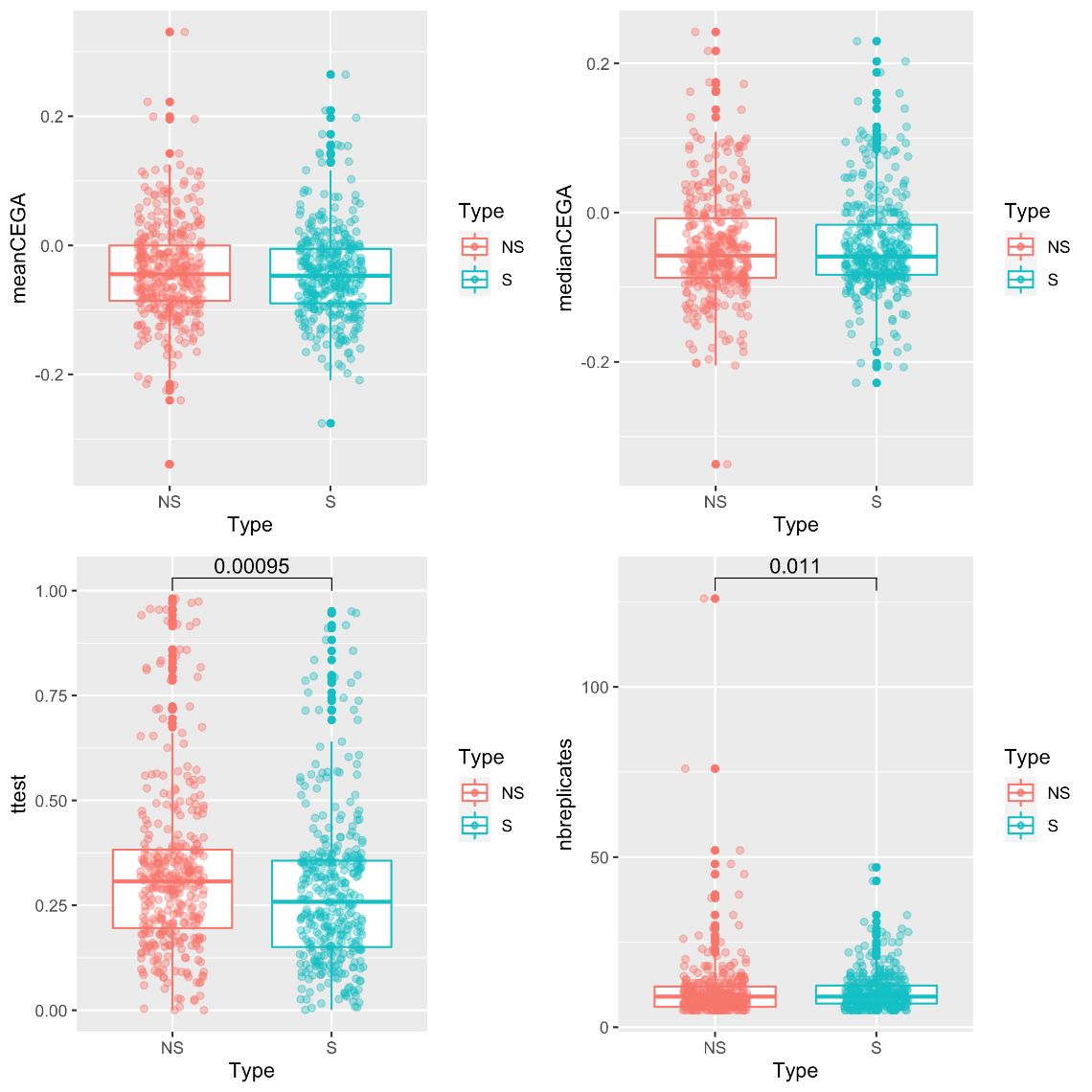

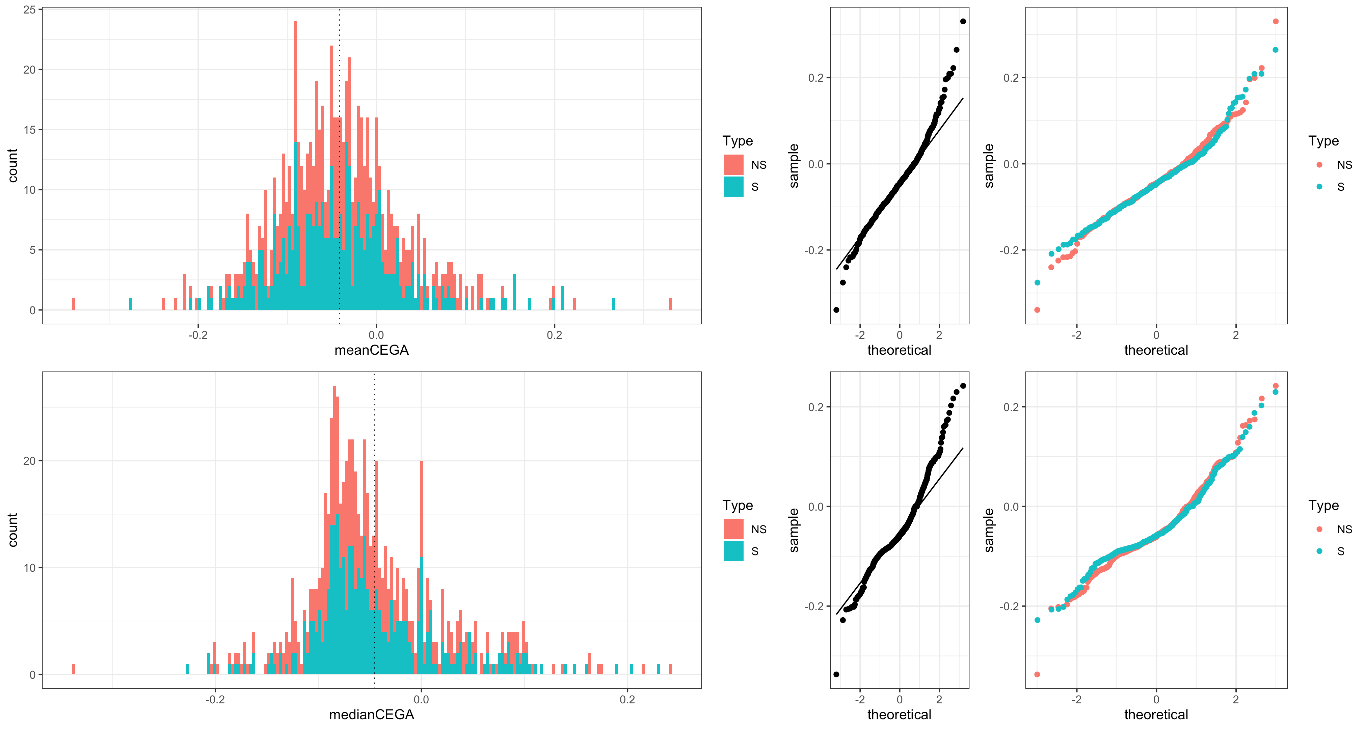

**Figure S7** Distribution of the mean and median CEGA values of all tested mutations and deletions according to their amino-acid change: synonymous or non-synonymous.

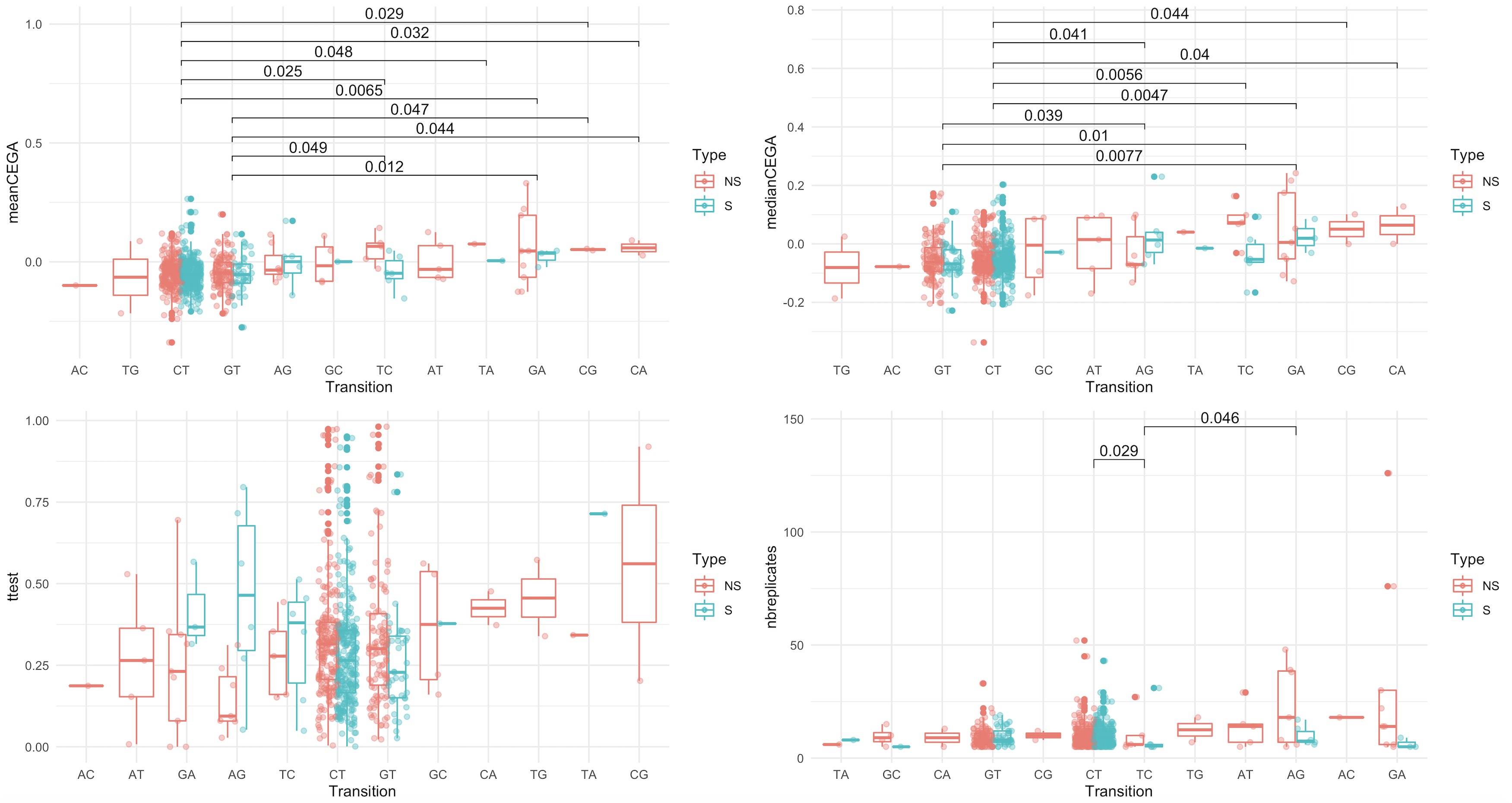

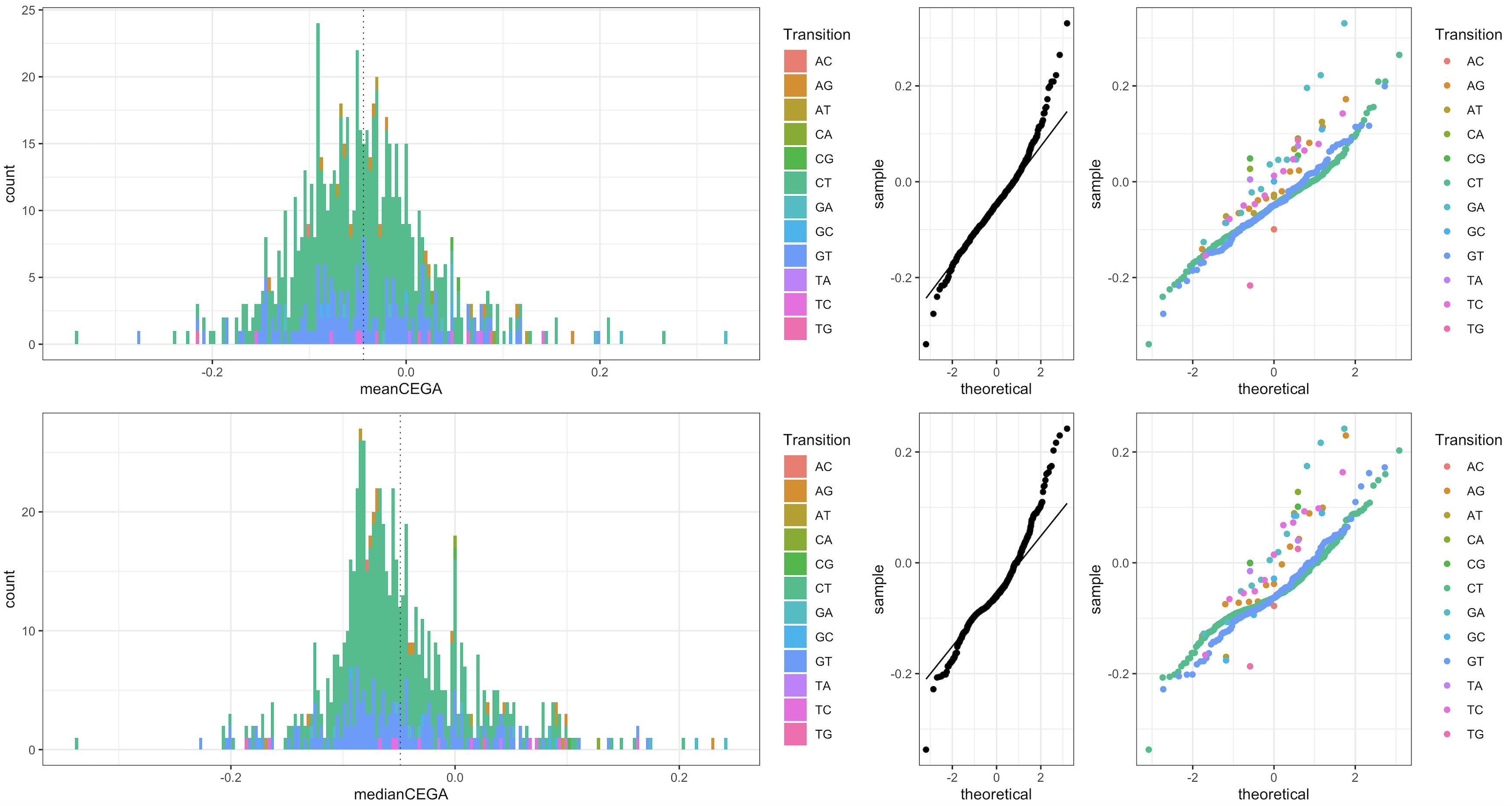

**Figure S8** Distribution of the mean and median CEGA values of all tested mutations and deletions according to their mutation type. Significant pairwise comparisons are highlighted at top. Mean and median CEGA values at C🡪T positions and G🡪T positions were the only transitions with inferred scores significantly less than 0 (one sample t-test *p*<2.2e-13, with no other transitions *p*<0.04).

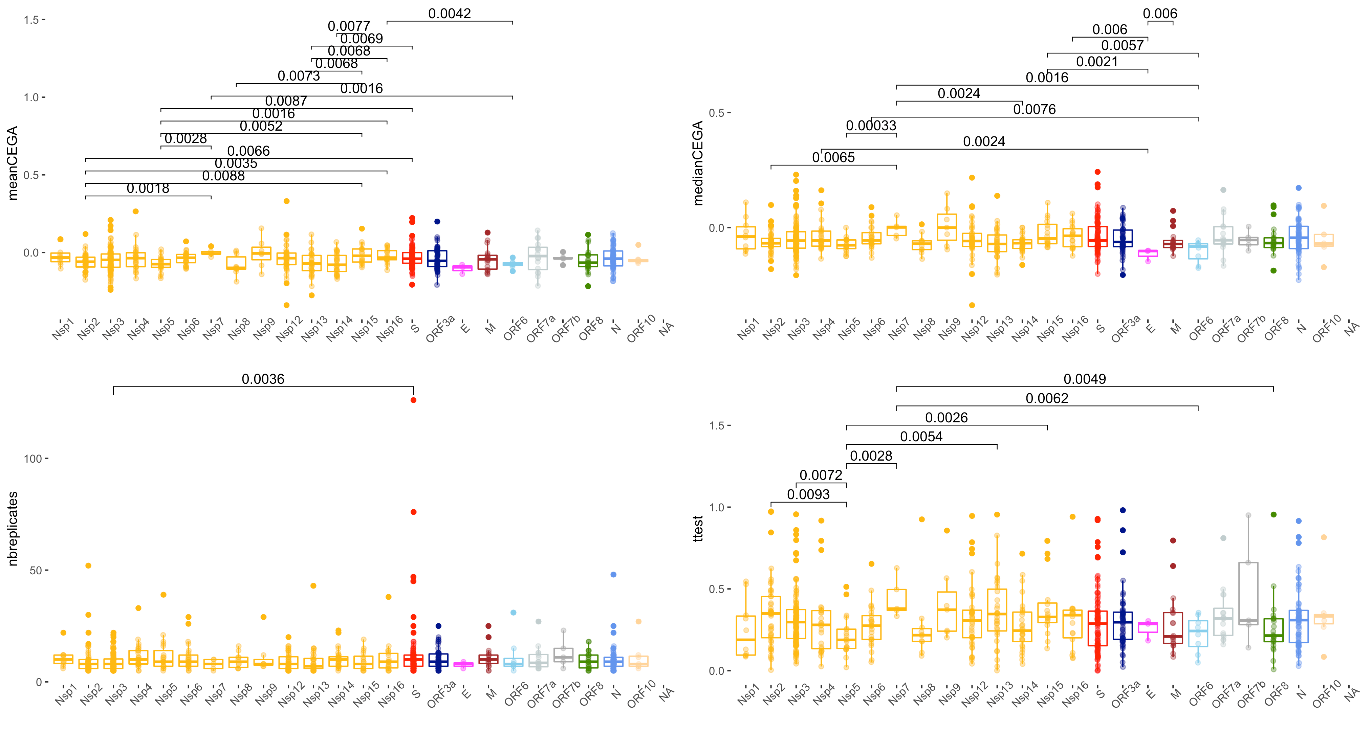

**Figure S9:** Boxplots providing the distributions associated to recurrent mutations at different genomic regions observed in the SARS-CoV-2 alignment. The top row provides the CEGA values estimated using the mean (left) and median (right) across nodes, while the bottom provides the significance and number of detected emergences (nbreplicates) and *p-*values of the paired t test on the number of offspring carrying vs. not carrying the mutation under scrutiny.

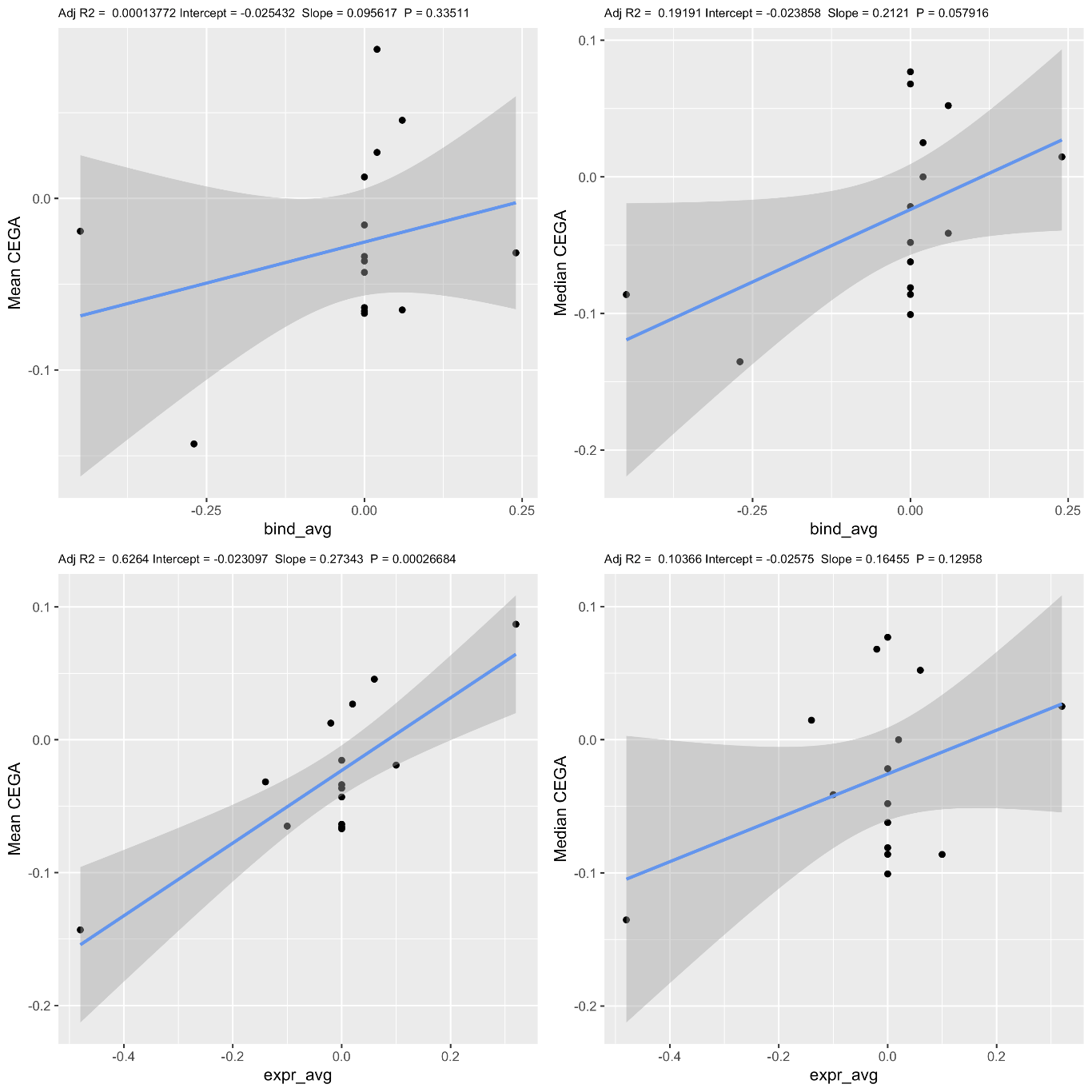

**Figure S10** Mean and median CEGA values and corresponding binding affinities and expression levels of a subset of seven mutations of the Receptor Binding Domain of the spike SARS-CoV-2 protein. Binding affinities and expression levels were computed by Starr et al. 2020.

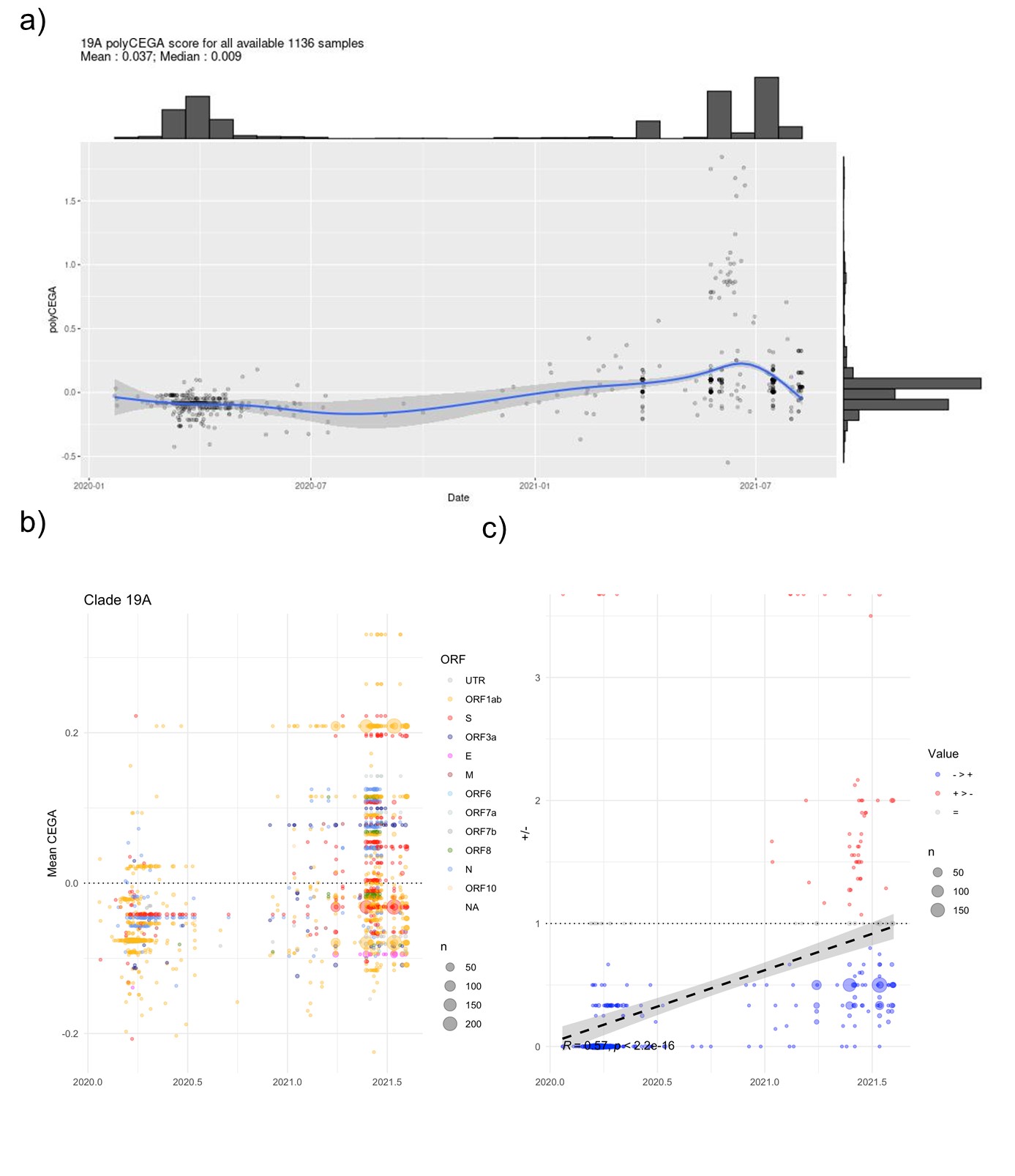

**Figure S11:** a) Poly-CEGA values for genome assemblies annotated to clade 19A. Each point represents an assembly of the clade under scrutiny and whose poly-CEGA value is read on the y axis and whose sampling date on the x-axis. The number of considered genome assemblies and mean and median poly-CEGA scores are noted at top left. b) Mutations being taken forward for the poly-CEGA computation over time (x-axis) coloured by major gene/ORF. c) Ratio of positive to negative CEGA scores per isolate through time. Line provides a simple linear regression with equation at bottom left.

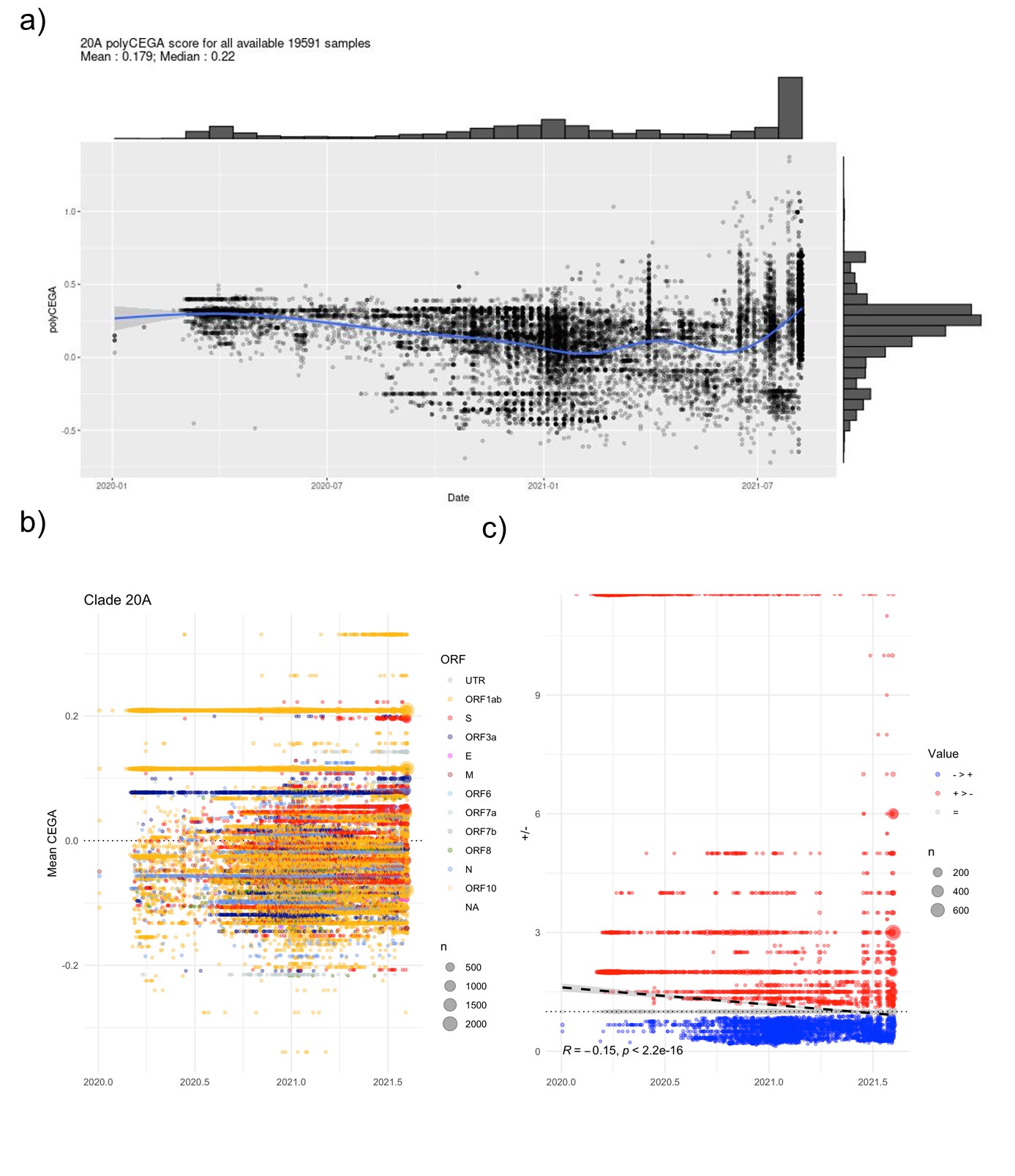

**Figure S12:** a) Poly-CEGA values for genome assemblies annotated to clade 20A. Each point represents an assembly of the clade under scrutiny and whose poly-CEGA value is read on the y axis and whose sampling date on the x-axis. The number of considered genome assemblies and mean and median poly-CEGA scores are noted at top left. b) Mutations being taken forward for the poly-CEGA computation over time (x-axis) coloured by major gene/ORF. c) Ratio of positive to negative CEGA scores per isolate through time. Line provides a simple linear regression with equation at bottom left.

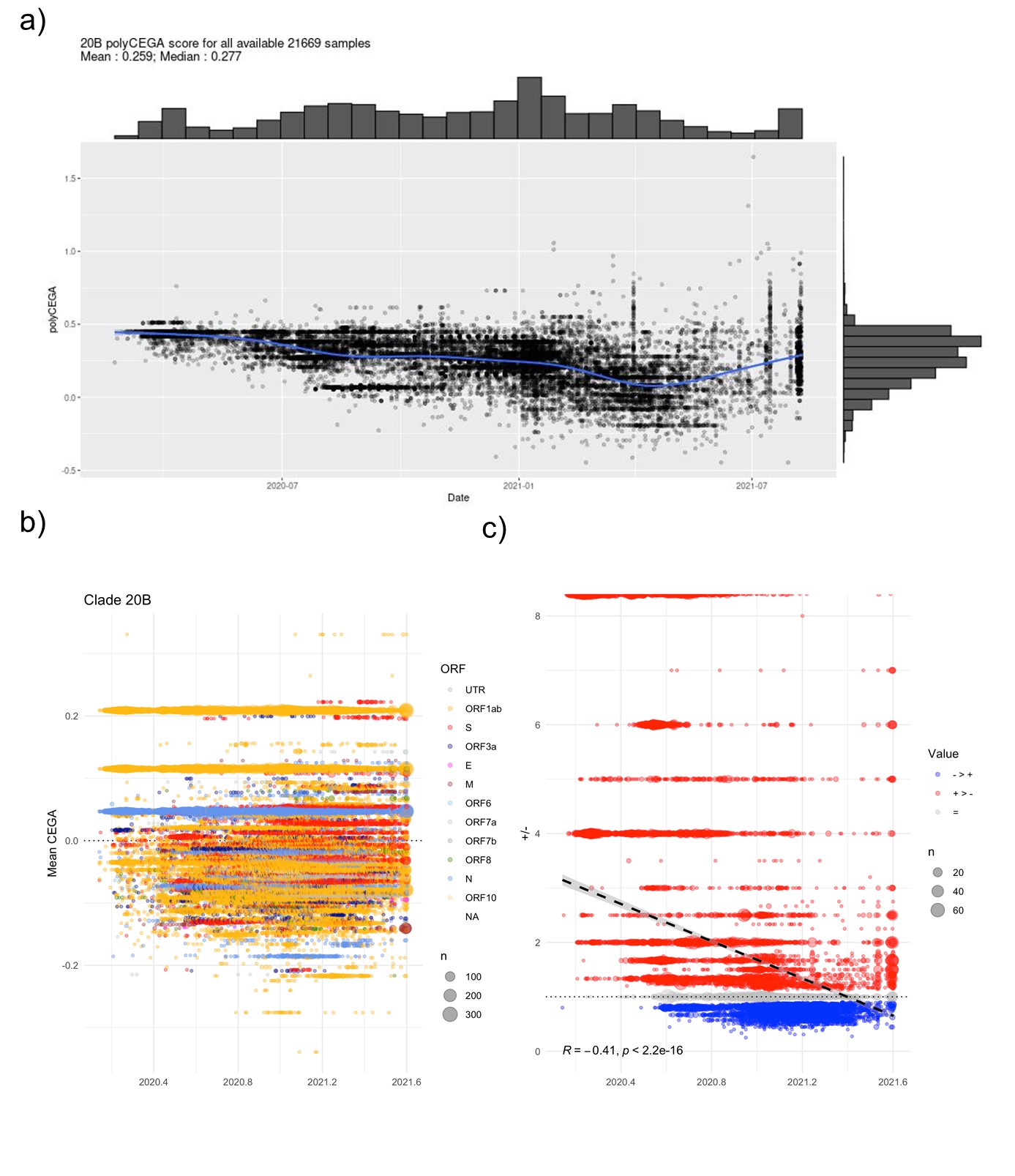

**Figure S13:** a) Poly-CEGA values for genome assemblies annotated to clade 20B. Each point represents an assembly of the clade under scrutiny and whose poly-CEGA value is read on the y axis and whose sampling date on the x-axis. The number of considered genome assemblies and mean and median poly-CEGA scores are noted at top left. b) Mutations being taken forward for the poly-CEGA computation over time (x-axis) coloured by major gene/ORF. c) Ratio of positive to negative CEGA scores per isolate through time. Line provides a simple linear regression with equation at bottom left.

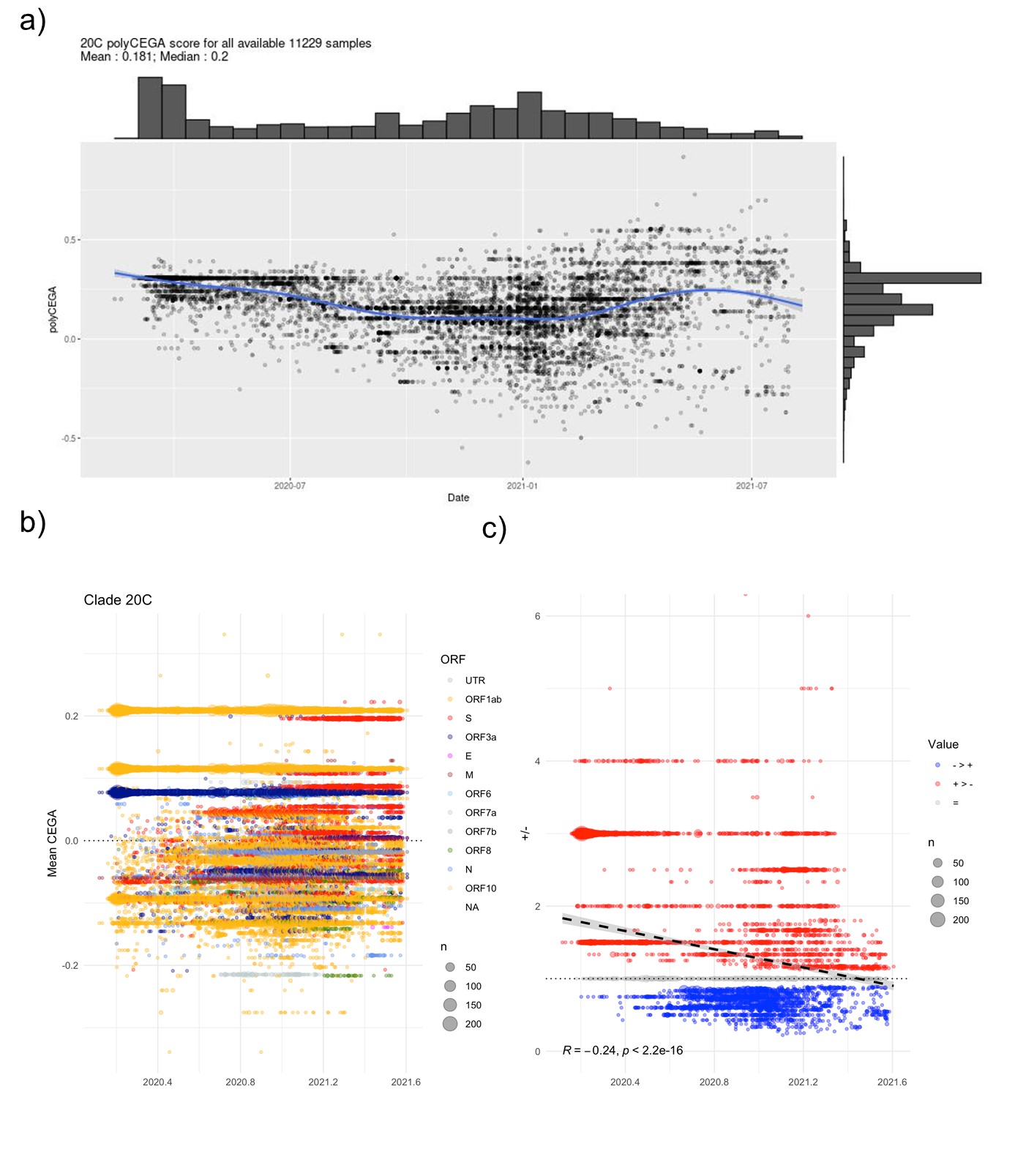

**Figure S14:** a) Poly-CEGA values for genome assemblies annotated to clade 20C. Each point represents an assembly of the clade under scrutiny and whose poly-CEGA value is read on the y axis and whose sampling date on the x-axis. The number of considered genome assemblies and mean and median poly-CEGA scores are noted at top left. b) Mutations being taken forward for the poly-CEGA computation over time (x-axis) coloured by major gene/ORF. c) Ratio of positive to negative CEGA scores per isolate through time. Line provides a simple linear regression with equation at bottom left.

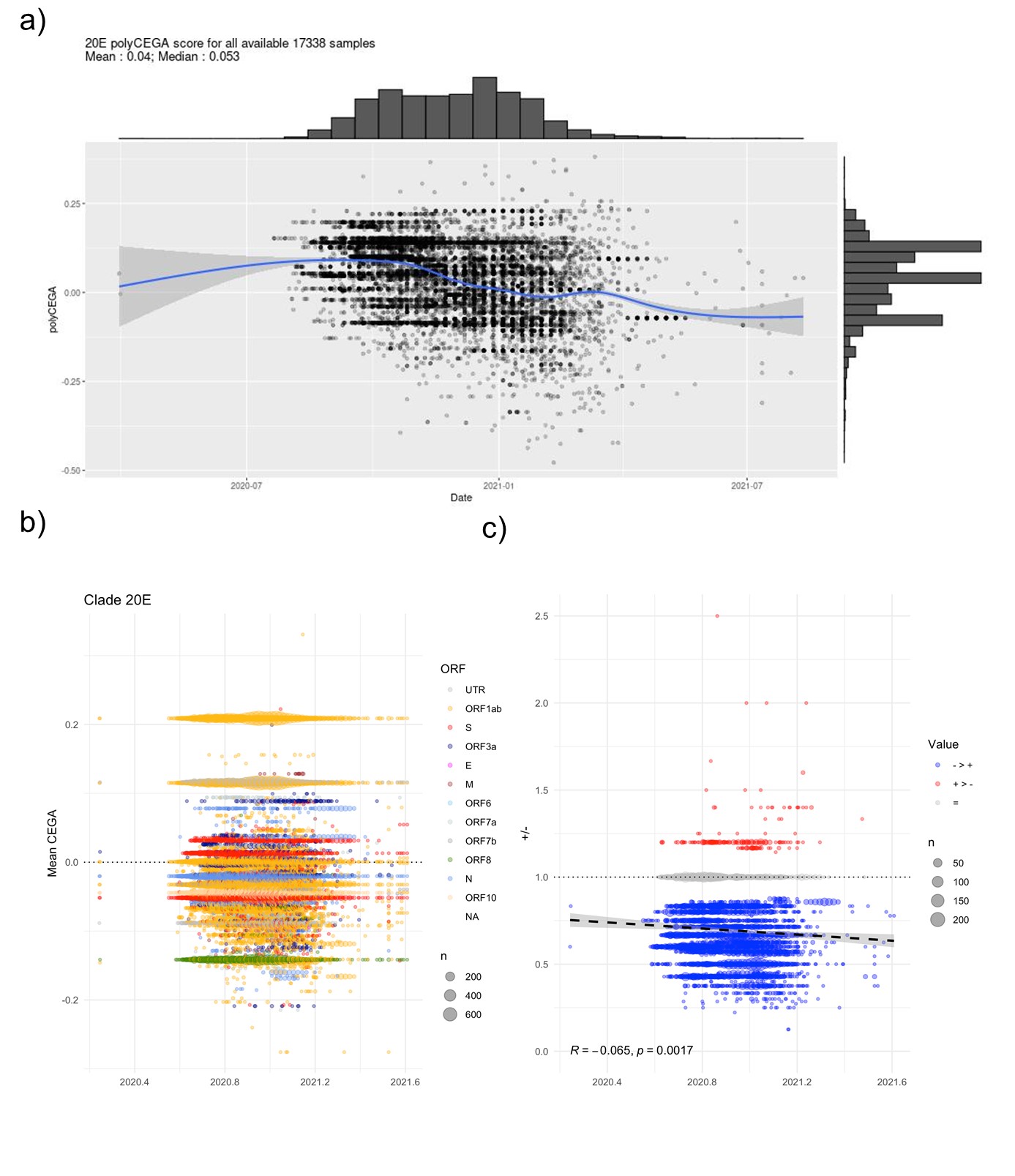

**Figure S15:** a) Poly-CEGA values for genome assemblies annotated to clade 20E. Each point represents an assembly of the clade under scrutiny and whose poly-CEGA value is read on the y axis and whose sampling date on the x-axis. The number of considered genome assemblies and mean and median poly-CEGA scores are noted at top left. b) Mutations being taken forward for the poly-CEGA computation over time (x-axis) coloured by major gene/ORF. c) Ratio of positive to negative CEGA scores per isolate through time. Line provides a simple linear regression with equation at bottom left.

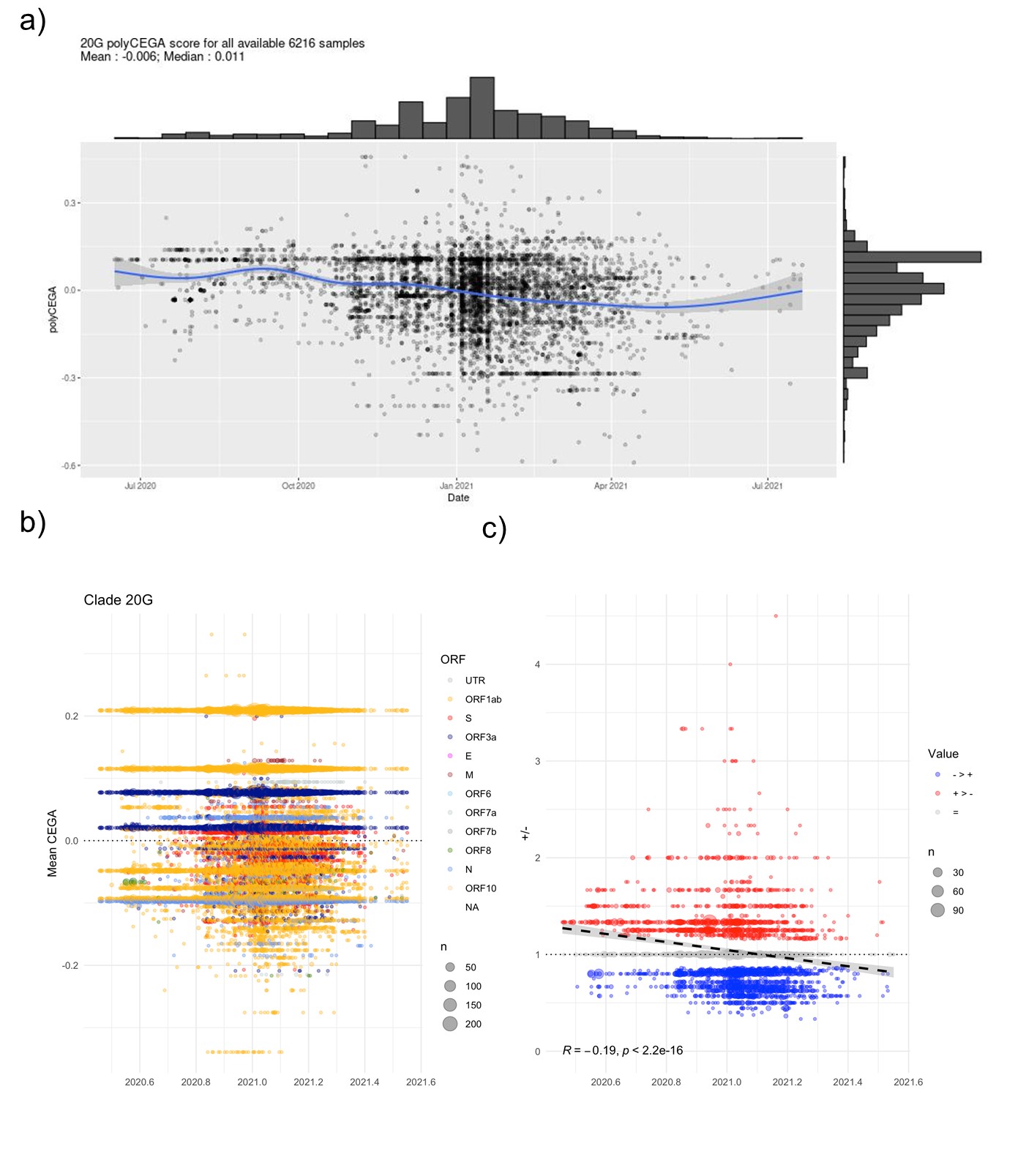

**Figure S16:** a) Poly-CEGA values for genome assemblies annotated to clade 20G. Each point represents an assembly of the clade under scrutiny and whose poly-CEGA value is read on the y axis and whose sampling date on the x-axis. The number of considered genome assemblies and mean and median poly-CEGA scores are noted at top left. b) Mutations being taken forward for the poly-CEGA computation over time (x-axis) coloured by major gene/ORF. c) Ratio of positive to negative CEGA scores per isolate through time. Line provides a simple linear regression with equation at bottom left.

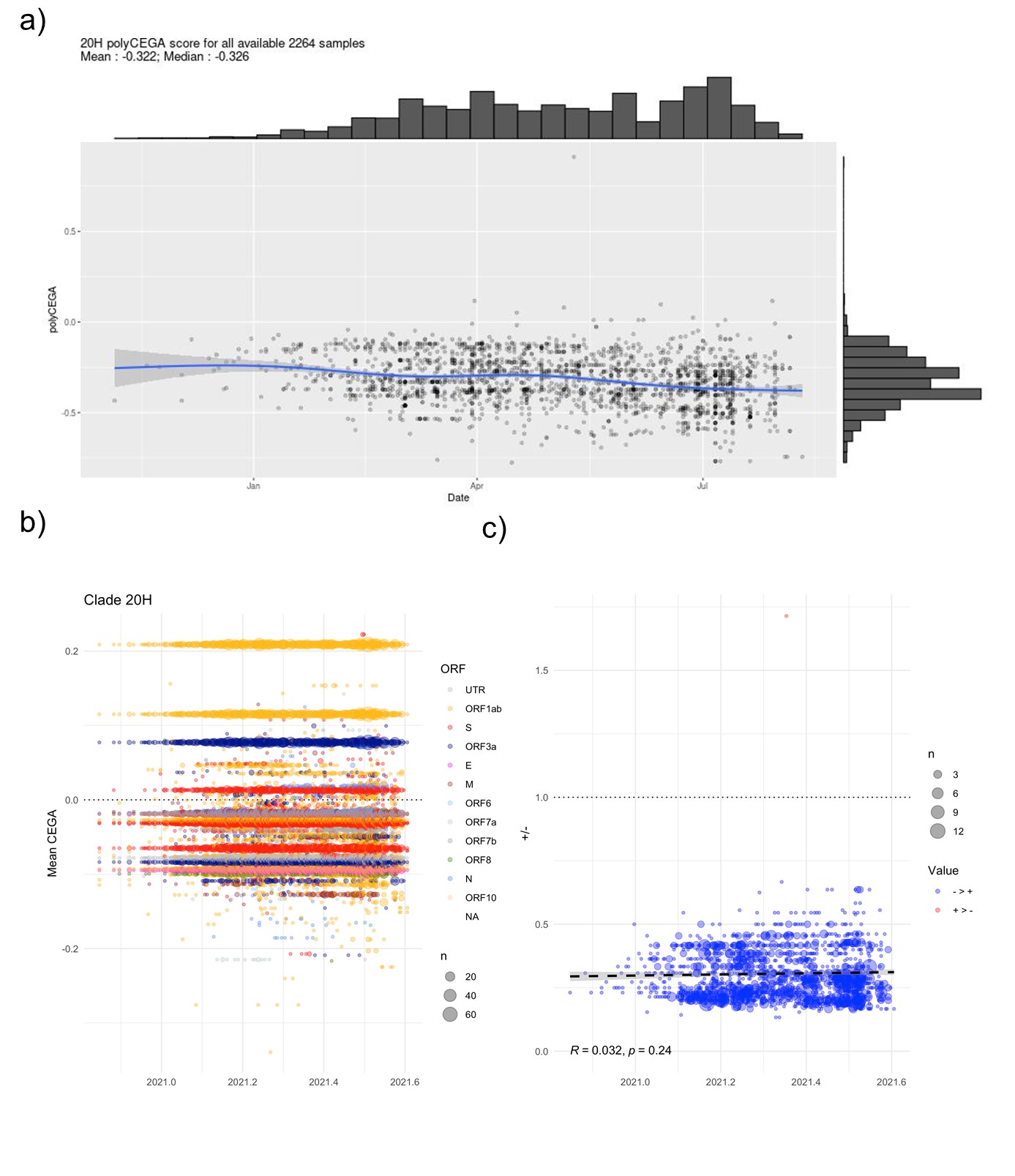

**Figure S17:** a) Poly-CEGA values for genome assemblies annotated to clade 20H. Each point represents an assembly of the clade under scrutiny and whose poly-CEGA value is read on the y axis and whose sampling date on the x-axis. The number of considered genome assemblies and mean and median poly-CEGA scores are noted at top left. b) Mutations being taken forward for the poly-CEGA computation over time (x-axis) coloured by major gene/ORF. c) Ratio of positive to negative CEGA scores per isolate through time. Line provides a simple linear regression with equation at bottom left.

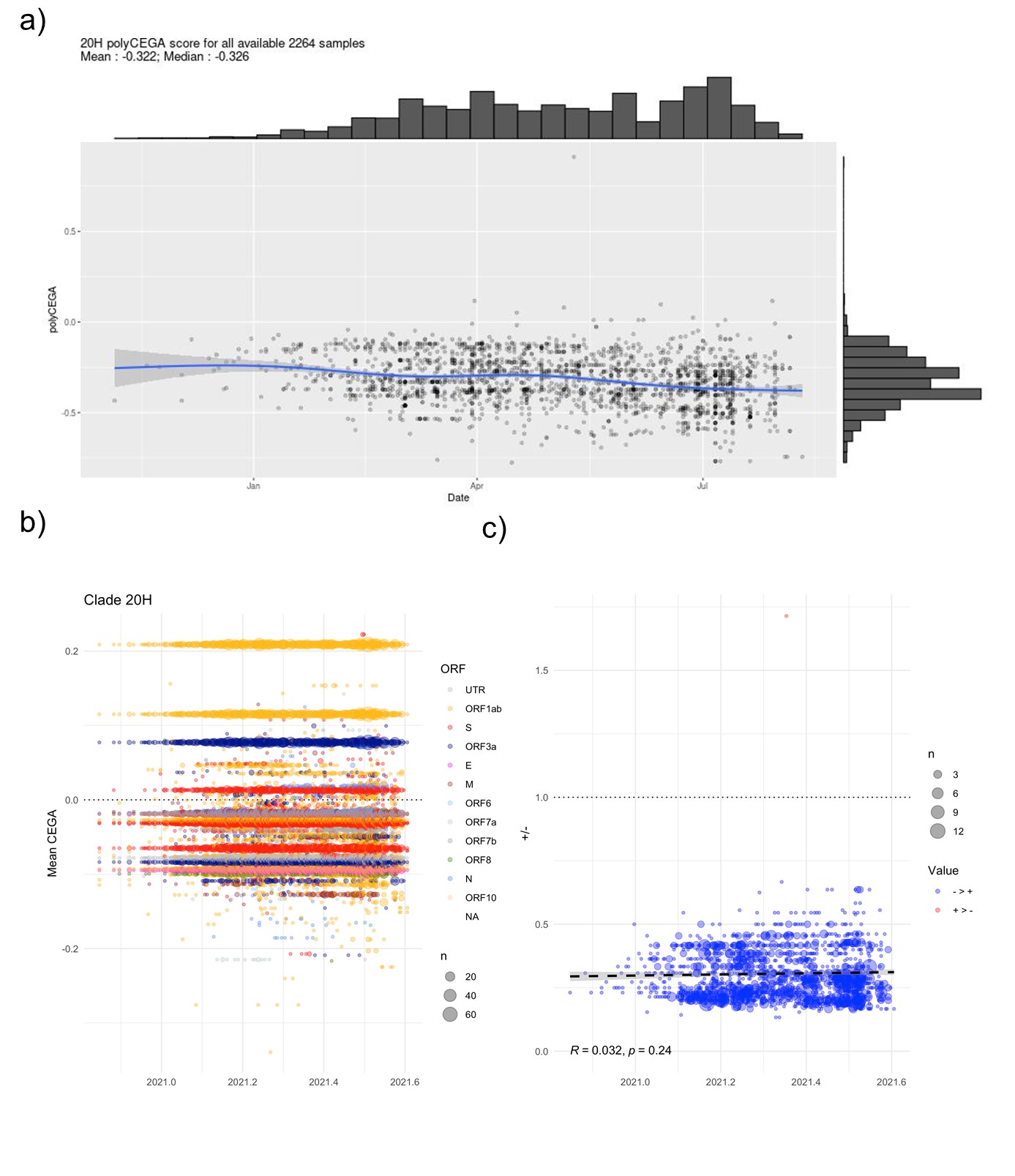

**Figure S18:** a) Poly-CEGA values for genome assemblies annotated to clade 20H. Each point represents an assembly of the clade under scrutiny and whose poly-CEGA value is read on the y axis and whose sampling date on the x-axis. The number of considered genome assemblies and mean and median poly-CEGA scores are noted at top left. b) Mutations being taken forward for the poly-CEGA computation over time (x-axis) coloured by major gene/ORF. c) Ratio of positive to negative CEGA scores per isolate through time. Line provides a simple linear regression with equation at bottom left.

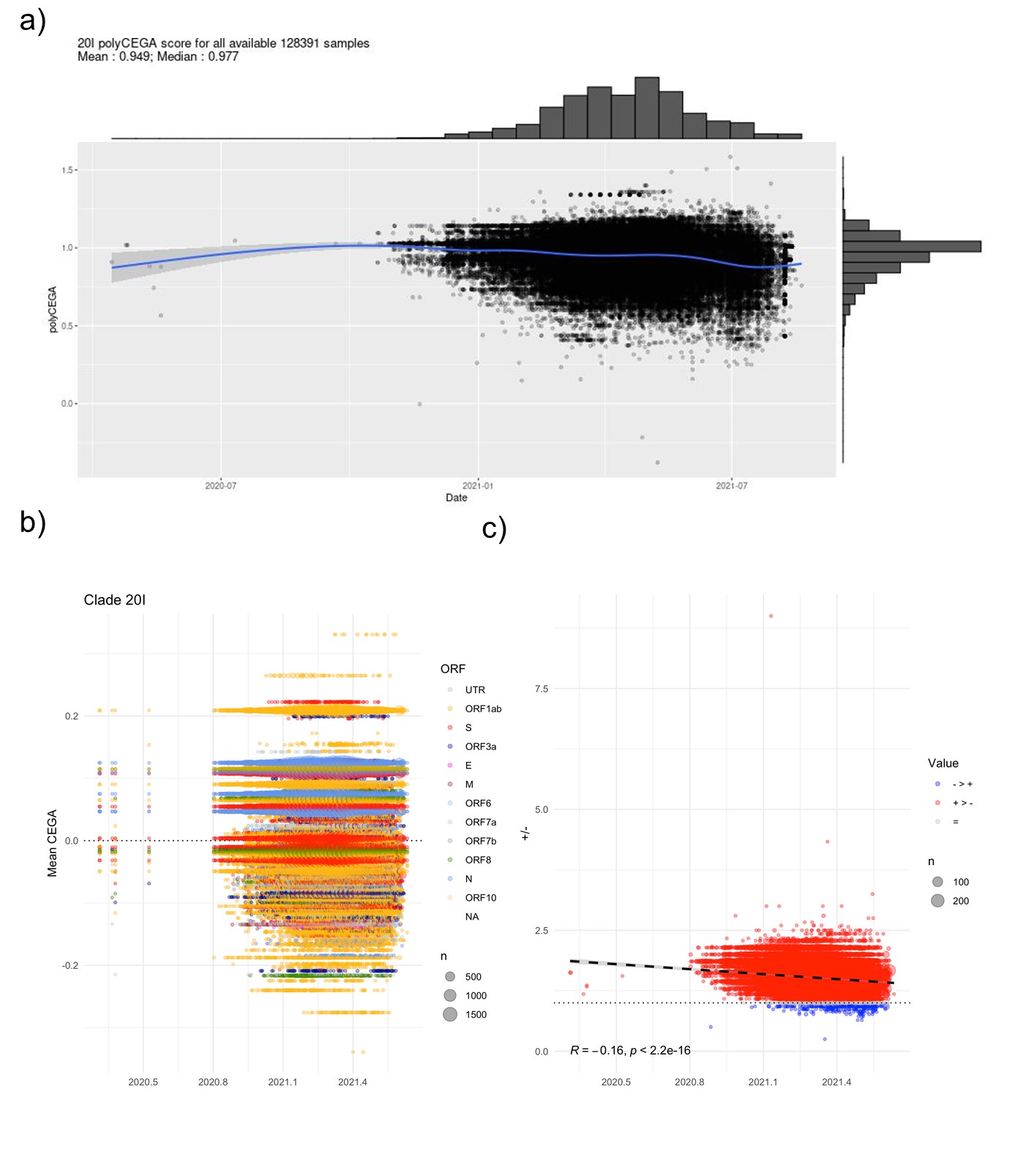

**Figure S19:** a) Poly-CEGA values for genome assemblies annotated to clade 20I based on a random sample of 80,000 assemblies. Each point represents an assembly of the clade under scrutiny and whose poly-CEGA value is read on the y axis and whose sampling date on the x-axis. The number of considered genome assemblies and mean and median poly-CEGA scores are noted at top left. b) Mutations being taken forward for the poly-CEGA computation over time (x-axis) coloured by major gene/ORF. c) Ratio of positive to negative CEGA scores per isolate through time. Line provides a simple linear regression with equation at bottom left.

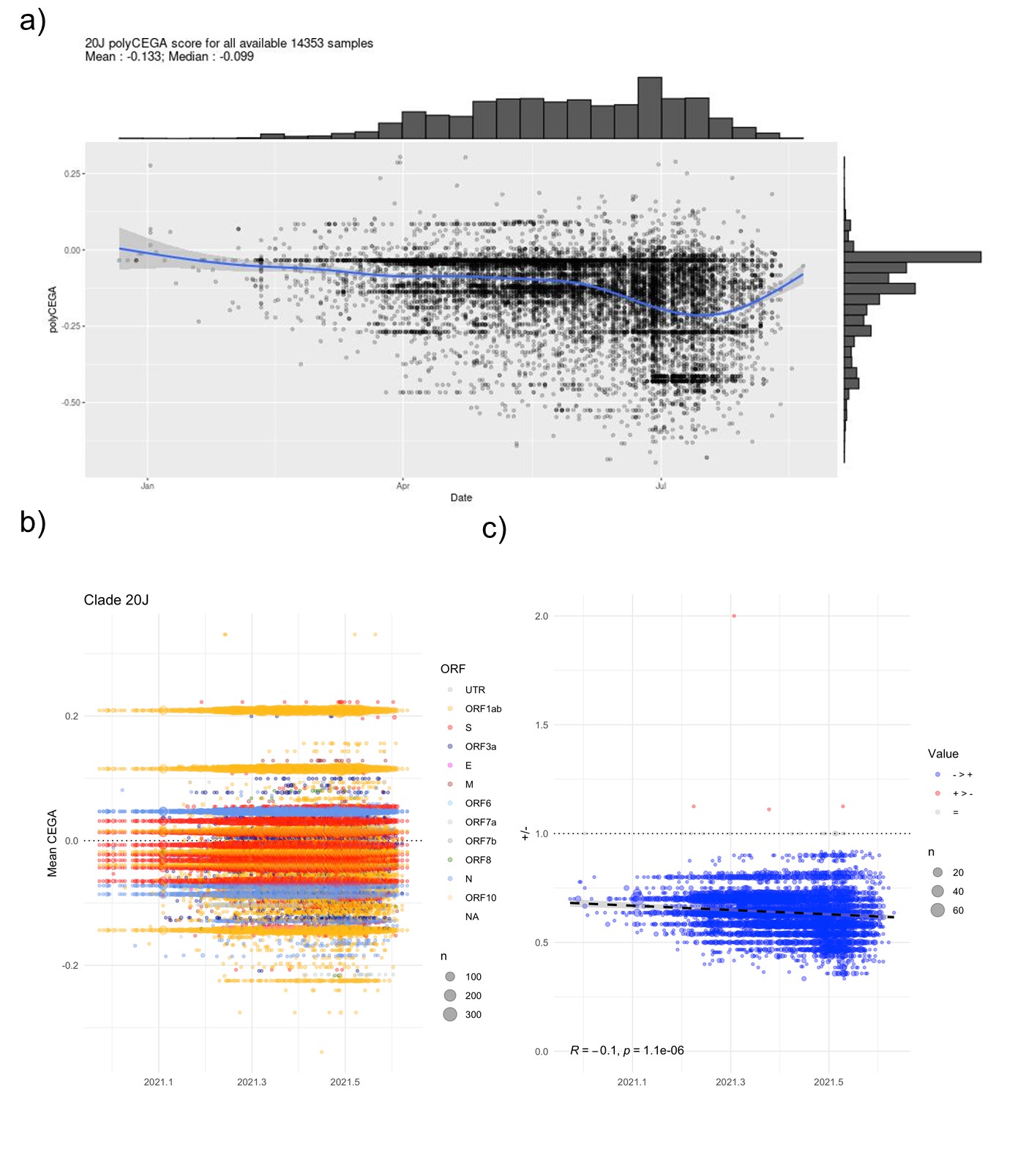

**Figure S20:** a) Poly-CEGA values for genome assemblies annotated to clade 20J. Each point represents an assembly of the clade under scrutiny and whose poly-CEGA value is read on the y axis and whose sampling date on the x-axis. The number of considered genome assemblies and mean and median poly-CEGA scores are noted at top left. b) Mutations being taken forward for the poly-CEGA computation over time (x-axis) coloured by major gene/ORF. c) Ratio of positive to negative CEGA scores per isolate through time. Line provides a simple linear regression with equation at bottom left.

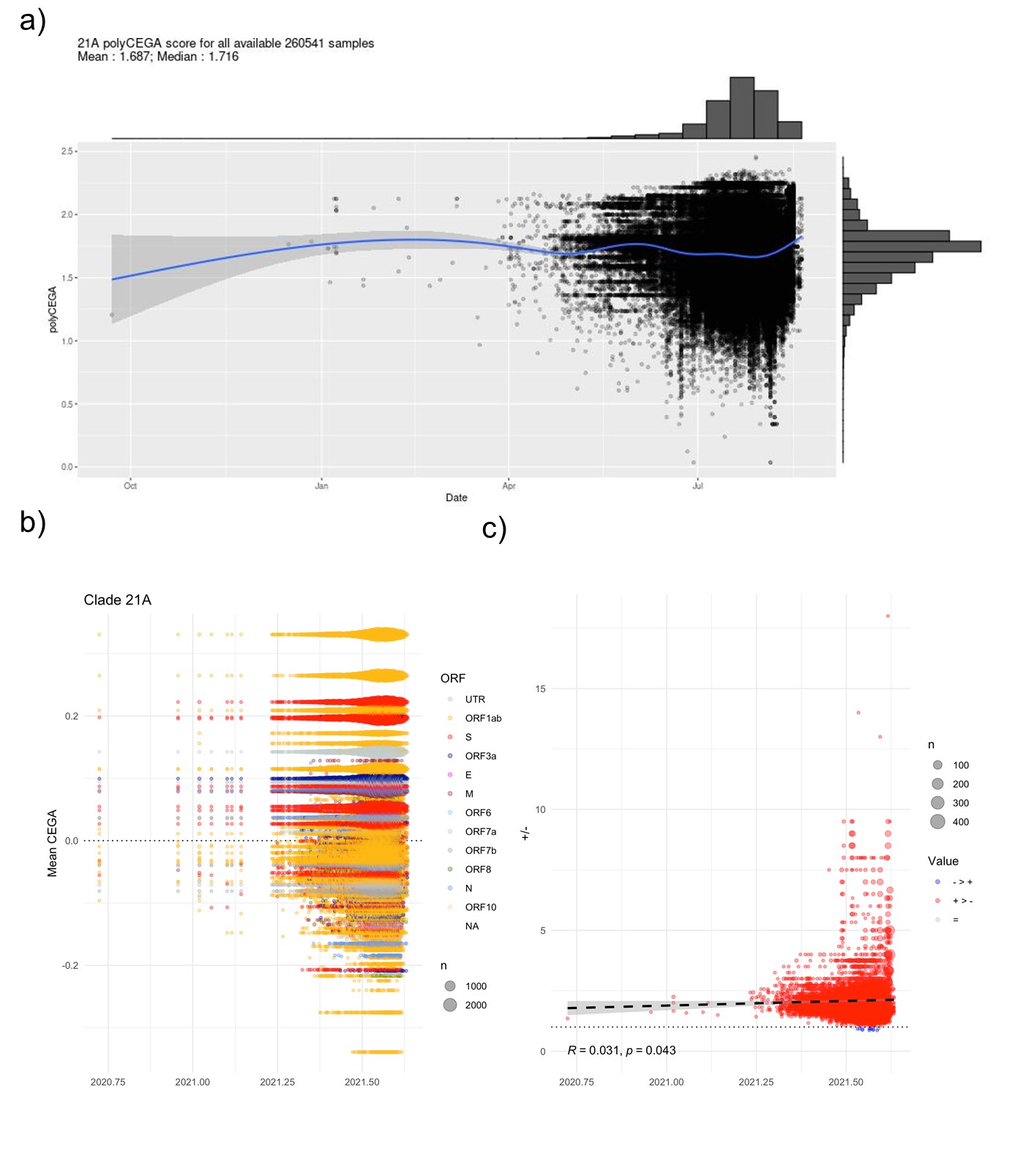

**Figure S21:** a) Poly-CEGA values for genome assemblies annotated to clade 21A based on a random sample of 80,000 assemblies. Each point represents an assembly of the clade under scrutiny and whose poly-CEGA value is read on the y axis and whose sampling date on the x-axis. The number of considered genome assemblies and mean and median poly-CEGA scores are noted at top left. b) Mutations being taken forward for the poly-CEGA computation over time (x-axis) coloured by major gene/ORF. c) Ratio of positive to negative CEGA scores per isolate through time. Line provides a simple linear regression with equation at bottom left.

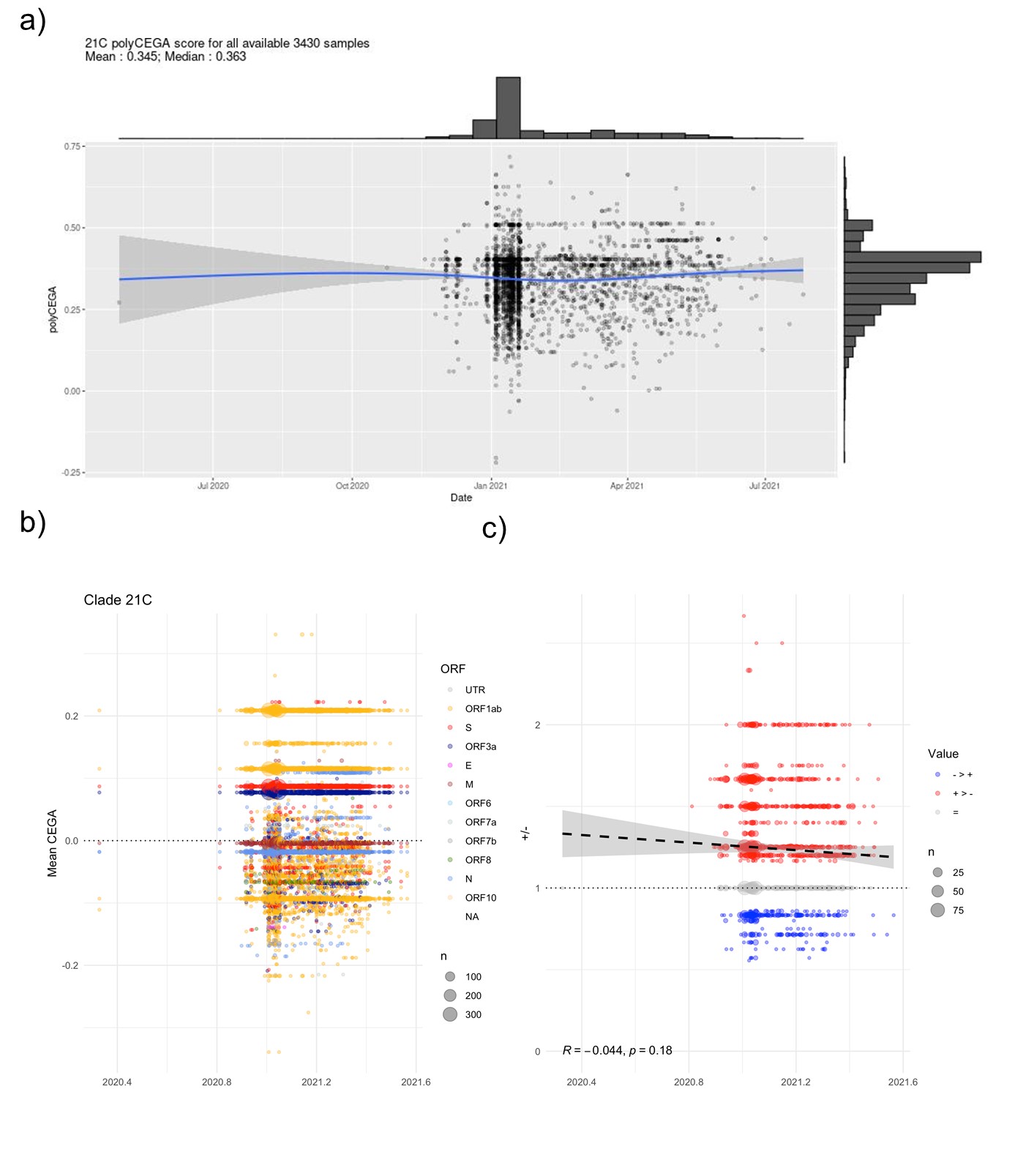

**Figure S22:** a) Poly-CEGA values for genome assemblies annotated to clade 21C. Each point represents an assembly of the clade under scrutiny and whose poly-CEGA value is read on the y axis and whose sampling date on the x-axis. The number of considered genome assemblies and mean and median poly-CEGA scores are noted at top left. b) Mutations being taken forward for the poly-CEGA computation over time (x-axis) coloured by major gene/ORF. c) Ratio of positive to negative CEGA scores per isolate through time. Line provides a simple linear regression with equation at bottom left.

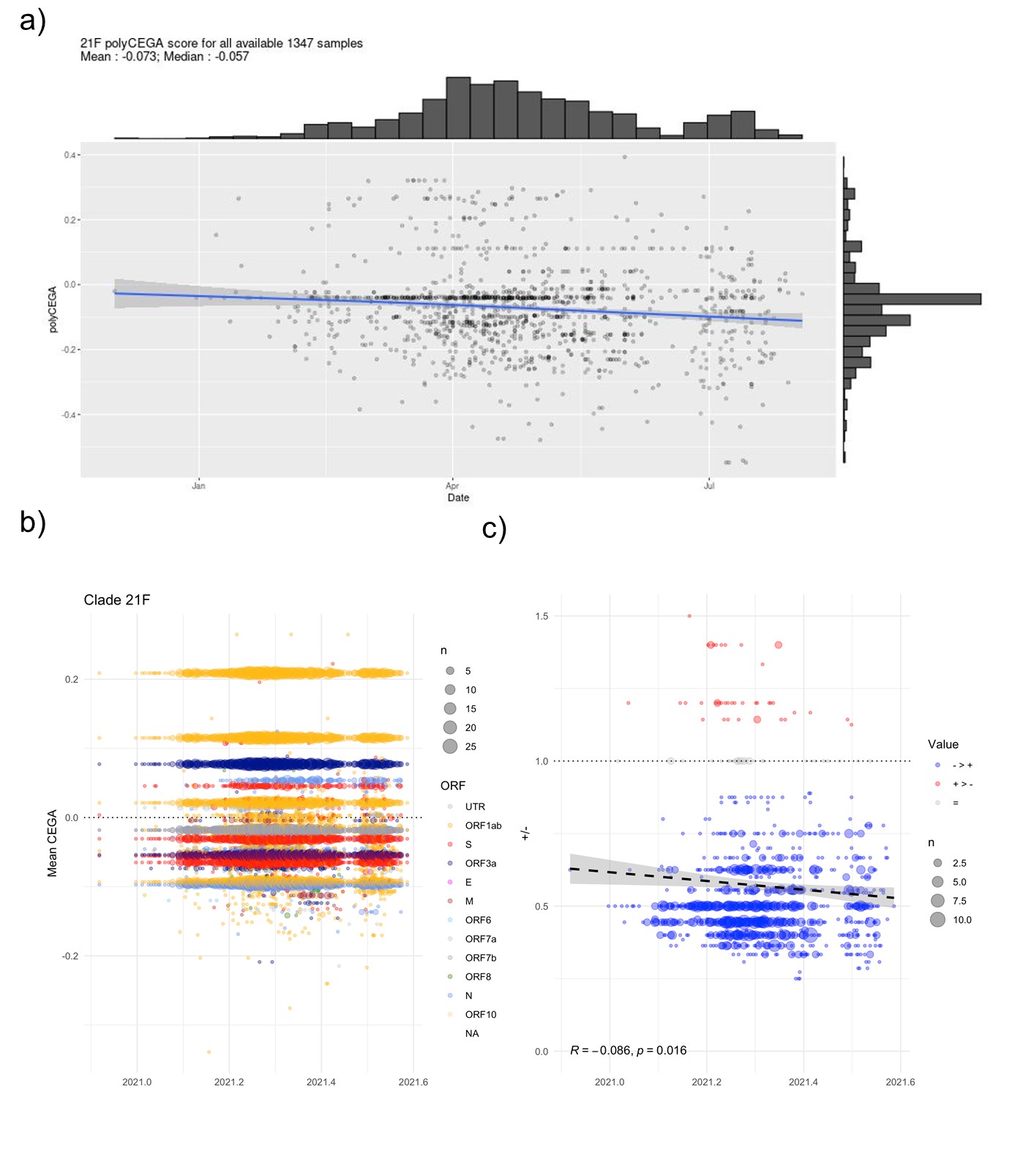

**Figure S23:** a) Poly-CEGA values for genome assemblies annotated to clade 21F. Each point represents an assembly of the clade under scrutiny and whose poly-CEGA value is read on the y axis and whose sampling date on the x-axis. The number of considered genome assemblies and mean and median poly-CEGA scores are noted at top left. b) Mutations being taken forward for the poly-CEGA computation over time (x-axis) coloured by major gene/ORF. c) Ratio of positive to negative CEGA scores per isolate through time. Line provides a simple linear regression with equation at bottom left.

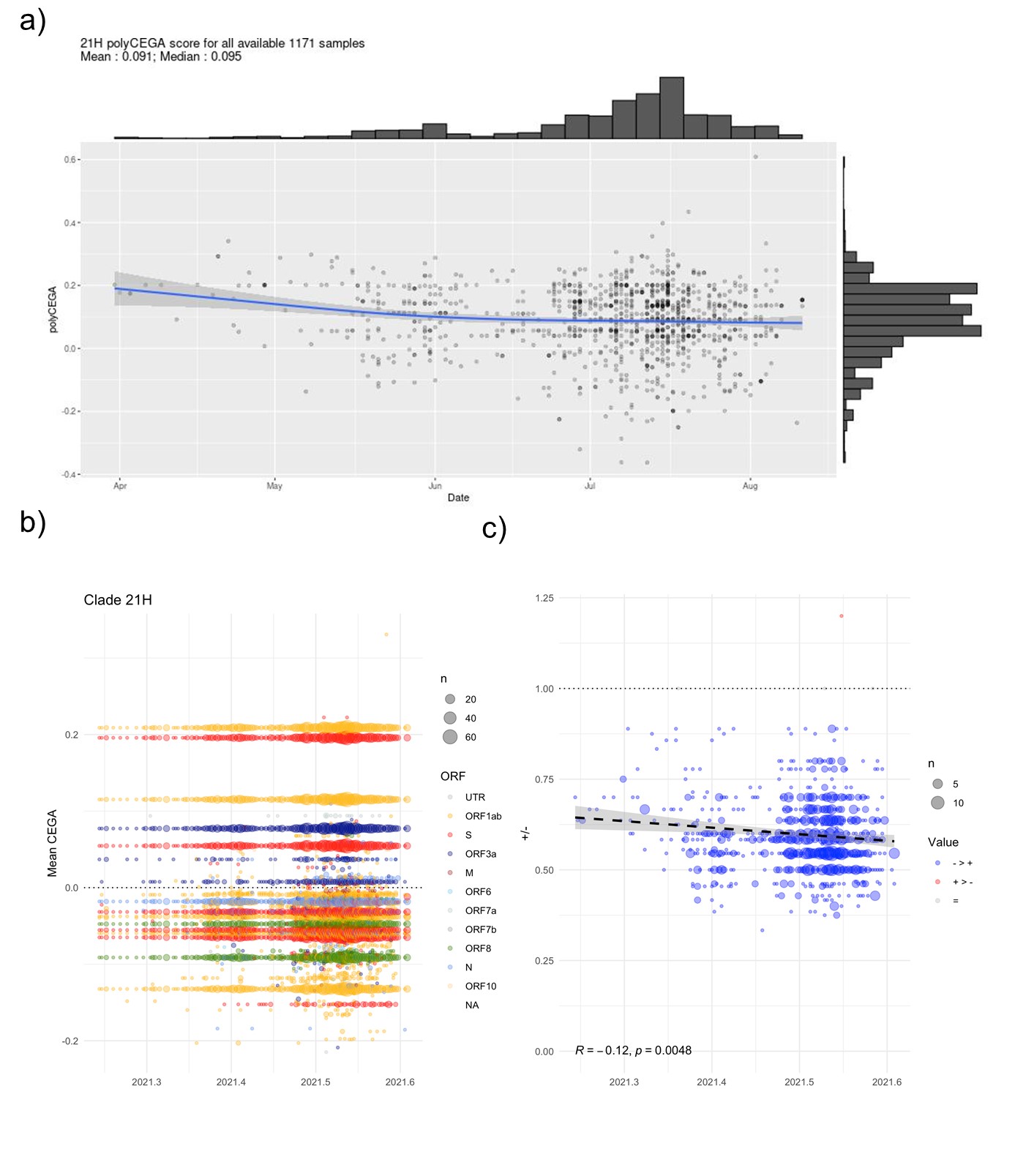

**Figure S24:** a) Poly-CEGA values for genome assemblies annotated to clade 21H. Each point represents an assembly of the clade under scrutiny and whose poly-CEGA value is read on the y axis and whose sampling date on the x-axis. The number of considered genome assemblies and mean and median poly-CEGA scores are noted at top left. b) Mutations being taken forward for the poly-CEGA computation over time (x-axis) coloured by major gene/ORF. c) Ratio of positive to negative CEGA scores per isolate through time. Line provides a simple linear regression with equation at bottom left.

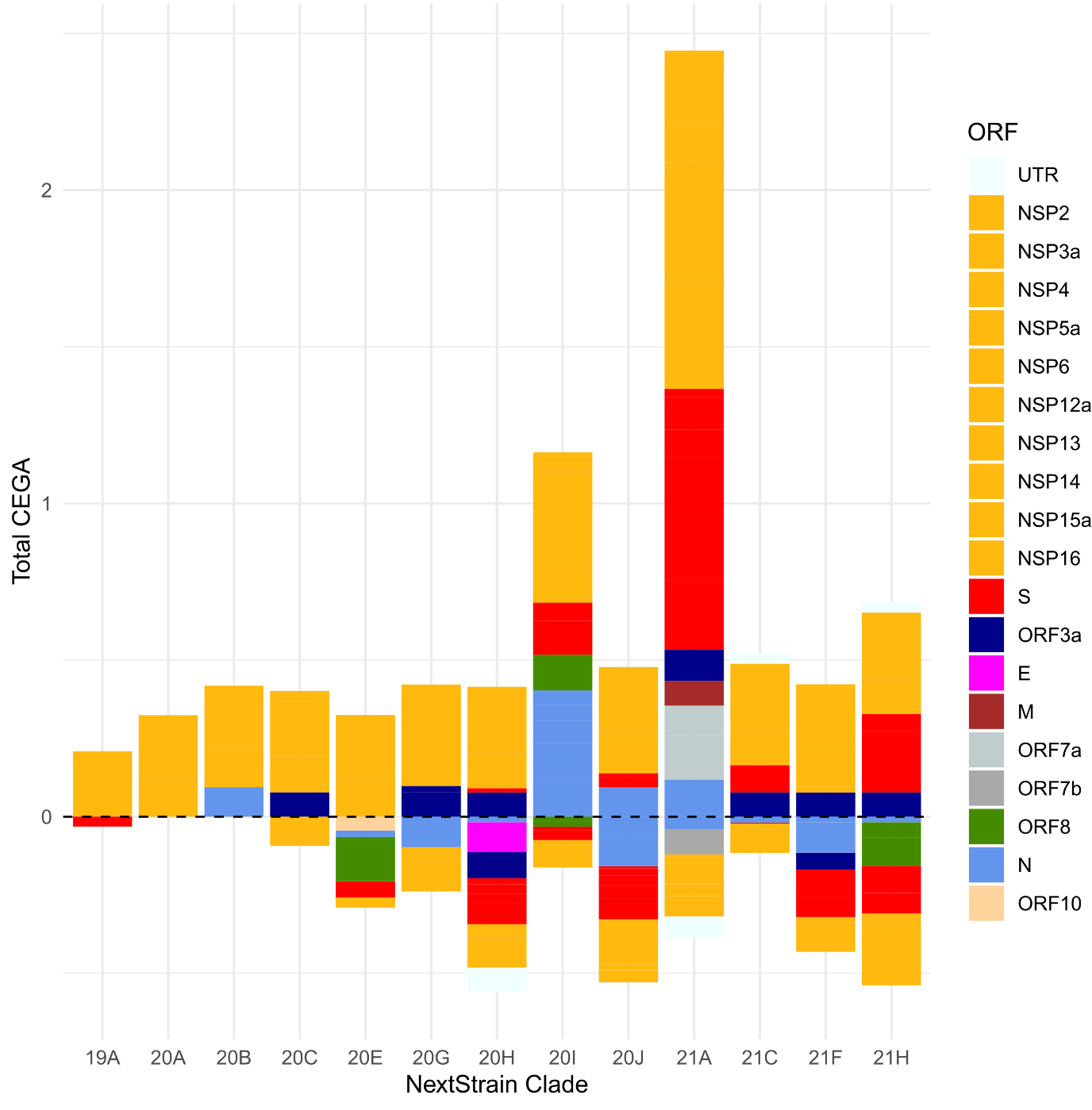

**Figure S25:** Contribution of mutations to polyCEGA scores estimated for 13 Nexstrain SARS-CoV-2 clades grouped by genomic region. For each clade, coloured blocks denote contributions of the mutations and deletions of each ORF to positive or negative CEGA scores. We only considered clades with >1,000 assemblies in our dataset and mutations with scores present in >50% of accessions in a clade.

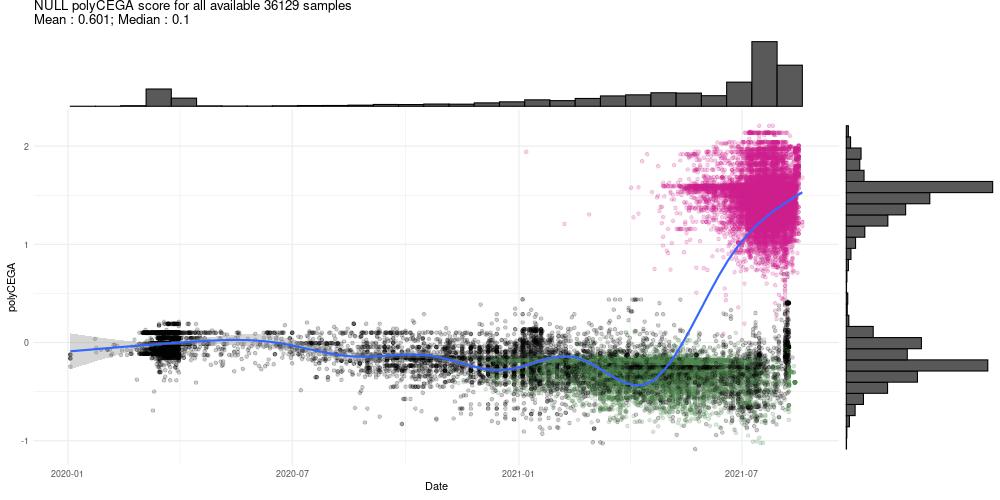

**Figure S26:** Poly-CEGA values (y-axis) computed from a dataset comprising all available assemblies from December 2019 to April 2020 and a random subset from April 2020 onwards, but with all assemblies assigned to the Alpha VoC removed (x-axis). Individual genomes (points) coloured in green label the Alpha VoC, those coloured in pink label the Delta VoC. All other considered genomes are coloured in black.

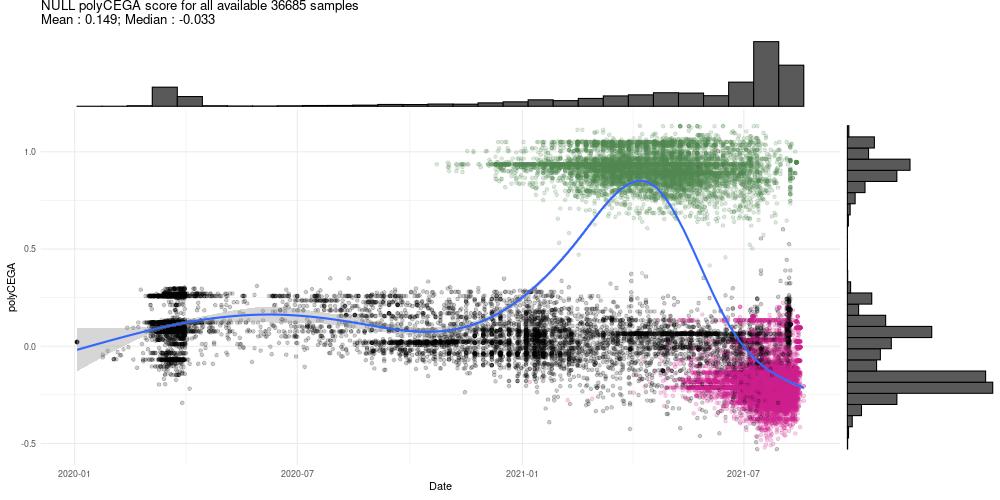

**Figure S27:** Poly-CEGA values (y-axis) computed from a dataset comprising all available assemblies from December 2019 to April 2020 and a random subset from April 2020 onwards, but with all assemblies assigned to the Delta VoC removed (x-axis). Individual genomes (points) coloured in green label the Alpha VoC, those coloured in pink label the Delta VoC. All other considered genomes are coloured in black.

**Table S1** Details of mutations and deletions defining VoCs. Orange denotes presence.

|  |  |  |  |  |  | 'Variant of Concern' | | | |
| --- | --- | --- | --- | --- | --- | --- | --- | --- | --- |
| **Protein name** | **Mutation** | **Nucleotide position** | **Nb. of emergences** | **Nb. of testable emergences** | **CEGA** | **Alpha (20I; 501Y.v1)** | **Beta (20H; 501Y.v2)** | **Gamma (20J; 501Y.v3)** | **Delta (21A)** |
| NSP6 | Δ106/108 | 11288-11296 | 724 | 28 | -0.022 |  |  |  |  |
| S | Δ69/70 | 21765-21770 | 550 | 14 | -0.010 |  |  |  |  |
| S | Δ156/157 | 22029-22034 | 1140 | 45 | 0.19 |  |  |  |  |
| S | K417T | 22812 | 102 | <5 | NA |  |  |  |  |
| S | K417N | 22813 | 106 | 7 | -0.019 |  |  |  |  |
| S | L452R | 22917 | 373 | 18 | 0.087 |  |  |  |  |
| S | E484K | 23012 | 321 | 22 | -0.065 |  |  |  |  |
| S | N501Y | 23063 | 494 | 29 | -0.032 |  |  |  |  |
| S | D614G | 23403 | 106 | <5 | NA |  |  |  |  |
| S | P681H | 23604 | NA | NA | NA |  |  |  |  |
| S | P681R | 23604 | 168 | 8 | 0.055 |  |  |  |  |

**Table S2.** External Excel document. SARS-CoV-2 homoplasies among 491,449 assemblies. No filtering has been applied apart from the masking of the first 150 and the last 300 nucleotides.

**Table S3.** External Excel document. Characteristics of the mutations and deletions tested using the CEGA scoring. In the case of deletions, we used the character "X" for alternate alleles.

**Table S4.** Coefficients of the linear models fitted to the polyCEGA scores of SARS-CoV-2 clades as of August 2021.

| **Nextrain Clade** | **WHO clade naming** | **Slope coefficient** | **p-val slope** | **Number of assemblies considered** |
| --- | --- | --- | --- | --- |
| 19A |  | 0,57 | <2.2e-16 | 1137 |
| 19B |  |  |  | 954 |
| 20A |  | -0,15 | <2.2e-16 | 19592 |
| 20B |  | -0,41 | <2.2e-16 | 21670 |
| 20C |  | -0,24 | <2.2e-16 | 11230 |
| 20D |  |  |  | 825 |
| 20E |  | -0,065 | 0,0017 | 17339 |
| 20F |  |  |  | 1 |
| 20G |  | -0,19 | <2.2e-16 | 6217 |
| 20H | Beta | 0,032 | 0,24 | 2265 |
| 20I | Alpha | -0,16 | <2.2e-16 | 80000* |
| 20J | Gamma | -0,1 | 1,10E-06 | 14353 |
| 21A | Delta | 0,031 | 0,043 | 80000* |
| 21B | Kappa |  |  | 411 |
| 21C | Epsilon | -0,044 | 0,18 | 3431 |
| 21D | Eta |  |  | 413 |
| 21E | Theta |  |  | 12 |
| 21F | Iota | -0,086 | 0,016 | 1348 |
| 21G | Lambda |  |  | 187 |
| 21H | Mu | -0,12 | 0,0048 | 1172 |
| *subsampled randomly to 80,000 accessions | | |  |  |
| Note clades with <1000 accessions were not considered | | | |  |

**Table S5.** External Excel document. GISAID metadata and originating and submitting laboratories.
